# Leptomeningeal anti-tumor immunity follows unique signaling principles

**DOI:** 10.1101/2023.03.17.533041

**Authors:** Jan Remsik, Xinran Tong, Russell Z. Kunes, Min Jun Li, Ahmed Osman, Kiana Chabot, Ugur T. Sener, Jessica A. Wilcox, Danielle Isakov, Jenna Snyder, Tejus A. Bale, Ronan Chaligné, Dana Pe’er, Adrienne Boire

## Abstract

Metastasis to the cerebrospinal fluid (CSF)-filled leptomeninges, or leptomeningeal metastasis (LM), represents a fatal complication of cancer. Proteomic and transcriptomic analyses of human CSF reveal a substantial inflammatory infiltrate in LM. We find the solute and immune composition of CSF in the setting of LM changes dramatically, with notable enrichment in IFN-γ signaling. To investigate the mechanistic relationships between immune cell signaling and cancer cells within the leptomeninges, we developed syngeneic lung, breast, and melanoma LM mouse models. Here we show that transgenic host mice, lacking IFN-γ or its receptor, fail to control LM growth. Overexpression of *Ifng* through a targeted AAV system controls cancer cell growth independent of adaptive immunity. Instead, leptomeningeal IFN-γ actively recruits and activates peripheral myeloid cells, generating a diverse spectrum of dendritic cell subsets. These migratory, CCR7+ dendritic cells orchestrate the influx, proliferation, and cytotoxic action of natural killer cells to control cancer cell growth in the leptomeninges. This work uncovers leptomeningeal-specific IFN-γ signaling and suggests a novel immune-therapeutic approach against tumors within this space.

## Main Text

Metastasis, or spread of cancer to distant anatomic sites, requires cancer cells to enter into and thrive within microenvironments unlike those of the primary tumor. In parallel, immune cells migrate throughout the organism and enter these same microenvironments as a counter-offensive, carrying out complex cellular tasks to control the growth of disseminated malignant cells. This balance may tip in favour of cancer cell growth for a variety of reasons, most simply when immune cells are excluded, as is the case for the majority of metastases to the central nervous system. An important exception to this rule is that of leptomeningeal metastasis (LM). The leptomeninges, the cerebrospinal fluid (CSF)-filled protective coverings, encase the central nervous system. Cancer cell entry into the leptomeningeal space, or LM, provokes a profound inflammatory response ^1–3^, clinically reminiscent of infectious meningitis ^4–6^. Despite this abundance of immune cells and intense inflammatory signals, leptomeningeal cancer cells persist and even thrive, a reflection of inflammation-mediated transcriptional changes within these cancer cells ^1, 4^. How and why these abundant infiltrating inflammatory cells fail to control cancer cell growth remains enigmatic. Previous work has uncovered non-canonical transcriptional and functional changes in macrophages in the setting of metastasis, suggesting that other immune cells within this anatomical compartment may also behave atypically ^4^. In addition, levels of inflammatory cytokines in the leptomeningeal space do not reflect those outside the leptomeninges, consistent with both intrathecal cytokine generation and alternative regulatory system(s) within this space ^7^. Formal investigations of this complex, anatomically site-specific, call-and-response between immune cells and cancer cells have remained incomplete, due to the lack of immunocompetent mouse models ^1, 8^ and to piecemeal computational approaches that do not encompass the entirety of cellular and humoral signalling within this under-appreciated anatomic space. To uncover the mechanisms whereby the immune system fails to control cancer cell growth in the leptomeningeal space, we comprehensively captured and molecularly dissected the immune response to LM in human disease and novel immunocompetent mouse models at both the cellular and humoral levels.

## Results

### Leptomeningeal metastasis generates an inflamed milieu

LM is a uniformly fatal neurologic consequence of cancer. Although any malignancy can result in LM, it most commonly results from breast, lung and melanoma primaries ^9^. Cancer cells enter into the leptomeningeal space accompanied by a host of leukocytes, as can be appreciated by classical techniques including CSF cytospins (**Fig.1 A**). To understand this leukocytic pleocytosis at a molecular level, we profiled CSF collected from breast and lung cancer patients with (n = 5) and without (n = 3) LM using the 10X platform, collecting single cell transcriptomes, assigning cell identity by clustering and marker gene expression (**Fig. 1B** **and fig. S1**). In the absence of LM, the CSF is nearly acellular and contains predominantly CD4+ T cells (**Fig. 1, A to C, and fig. S1**). Pathological processes that do not directly involve the leptomeninges, such as parenchymal or dural metastases, remodel the CSF cell landscape towards reactive myeloid cells (**fig. S1, D to F**). CSF from patients harboring LM was pleocytic and contained cells from a spectrum of lymphoid and myeloid lineages. To assess the molecular hallmarks of CSF pleocytosis and capture cell-to-cell communication, we subjected CSF from patients with breast cancer, lung cancer, and melanoma primaries, with and without LM, to targeted proteomic analysis by proximity extension assay (**fig. S2, A to C, and tables S1 to 3**) ^9^. In the presence of LM, CSF demonstrated a robust influx of soluble inflammatory ligands; 15 of these molecules were shared across the three tumor types (**fig. S2D**). Extending this cohort to include patients with a wide variety of solid tumor primaries confirmed elevated CSF levels of IFN-**γ** in the presence of LM (**Fig. 1D**). Moreover, elevated CSF IFN-**γ** levels at diagnosis were associated with improved overall survival (**Fig. 1E**). IFN-**γ** is well-known to exhibit both pro-tumorigenic and tumor-supressive actions in a context-dependent manner; the presence of inflammatory and anti-inflammatory signals in cancer-infiltrated leptomeninges suggested a dense signaling network not clearly consistent with monotone behavior and canonical pathways. We therefore pursued formal identification of downstream leptomeningeal effectors of IFN-**γ** and the functional consequence(s) of their activation.

**Fig. 1.**
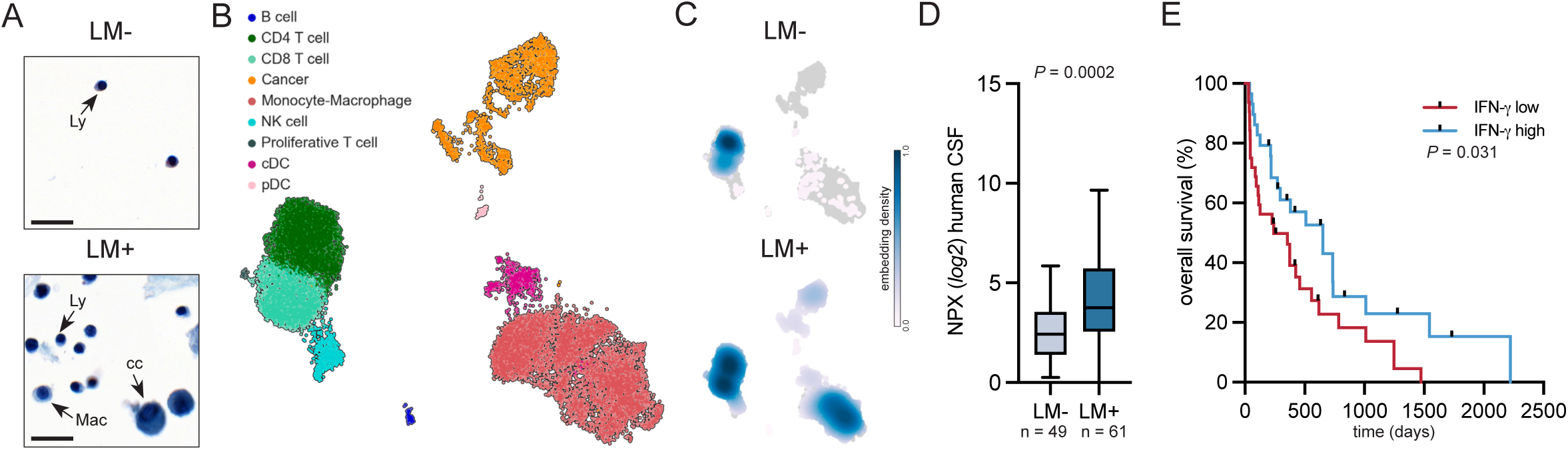
Inflammation-induced pleocytosis in patients with leptomeningeal metastasis. (**A**) Representative images of Giemsa-stained cytospins from cancer patients without (top) and with leptomeningeal metastasis (LM, bottom) with major cell populations indicated as Ly - lymphocyte, Mac - monocyte-macrophage, cc - cancer cell (n = 5 *per* group, scale bar = 20 μm). (**B**) UMAP projection of human CSF immune cell types and cancer cells, isolated from cancer patients without (n = 3 patients and n = 1,196 cells) and with (n = 5 patients and n = 16,022 cells) LM. LM+ samples were retrieved from GSE150660 and colored by cell type ^4^. See also fig. S1 and Methods for experimental overview, cell type annotations, and quality control plots. (**C**) Embedding density plots from LM- and LM+ patients, showing relative cell type abundance *per* condition, projected onto UMAP. (**D**) Relative CSF IFN-**γ** levels in cancer patients with or without LM from wide array of solid tumors, as determined by proximity extension assay (LM- n = 49, LM+ n = 61). NPX - normalized protein expression. See also fig. S2. (**E**) Kaplan-Meier plot showing, post-LM diagnosis survival in relation to CSF IFN-**γ** levels at diagnosis. Logrank test (IFN-**γ** high n = 29; IFN-**γ** low n = 32; cut-off NPX = 4).

### Interferon-**γ** regulates leptomeningeal metastatic growth

To enable these mechanistic studies, we leveraged iterative *in vivo* selection to generate six immunocompetent mouse LeptoM lines on two genetic backgrounds (**fig. S3A**) ^1^. These cell lines, subpopulations of the founding parental line, are phenotypically and transcriptomically distinct from their parental or brain parenchyma-tropic counterparts (**fig. S3, B to M**). Moreover, these LeptoM models faithfully recapitulate key histological and oncological features of human LM including CSF pleocytosis (**Fig. 2A**) and brisk pace of illness (**fig. S3, B to M**). To capture the complexity of leptomeningeal immune infiltrate at systems level, we performed proteogenomic analysis of mouse leptomeningeal immune infiltrate with 198 barcoded antibodies targeting cell surface epitopes and non-targeting isotype controls, coupled with single cell RNA-sequencing on the 10X platform; single-cell CITE-seq ^10^. This approach enables granular identification of immune cell subtypes and their origin. The mouse models mimic human CSF cellular composition in the setting of LM: a dramatic influx of leukocytes is observed, evenly split between myeloid and lymphoid populations (**Fig. 2, B and C, and fig. S4**, compare **Fig. 1** **B and C, and fig S1E**). In both human and mouse LM, the myeloid compartment is populated by monocyte-macrophages, and to a lesser extent, dendritic cells (DCs) (**fig. S1E, and fig S4**). In the absence of LM, normal human CSF is T-cell predominant, whereas disease-free mouse leptomeninges are populated by B cells and neutrophils. Despite species-specific differences in the in the absence of malignancy, the presence of LM drives the CSF cellular composition to a common, myeloid-dominant pleocytosis, independent of vertebrate host.

**Fig. 2.**
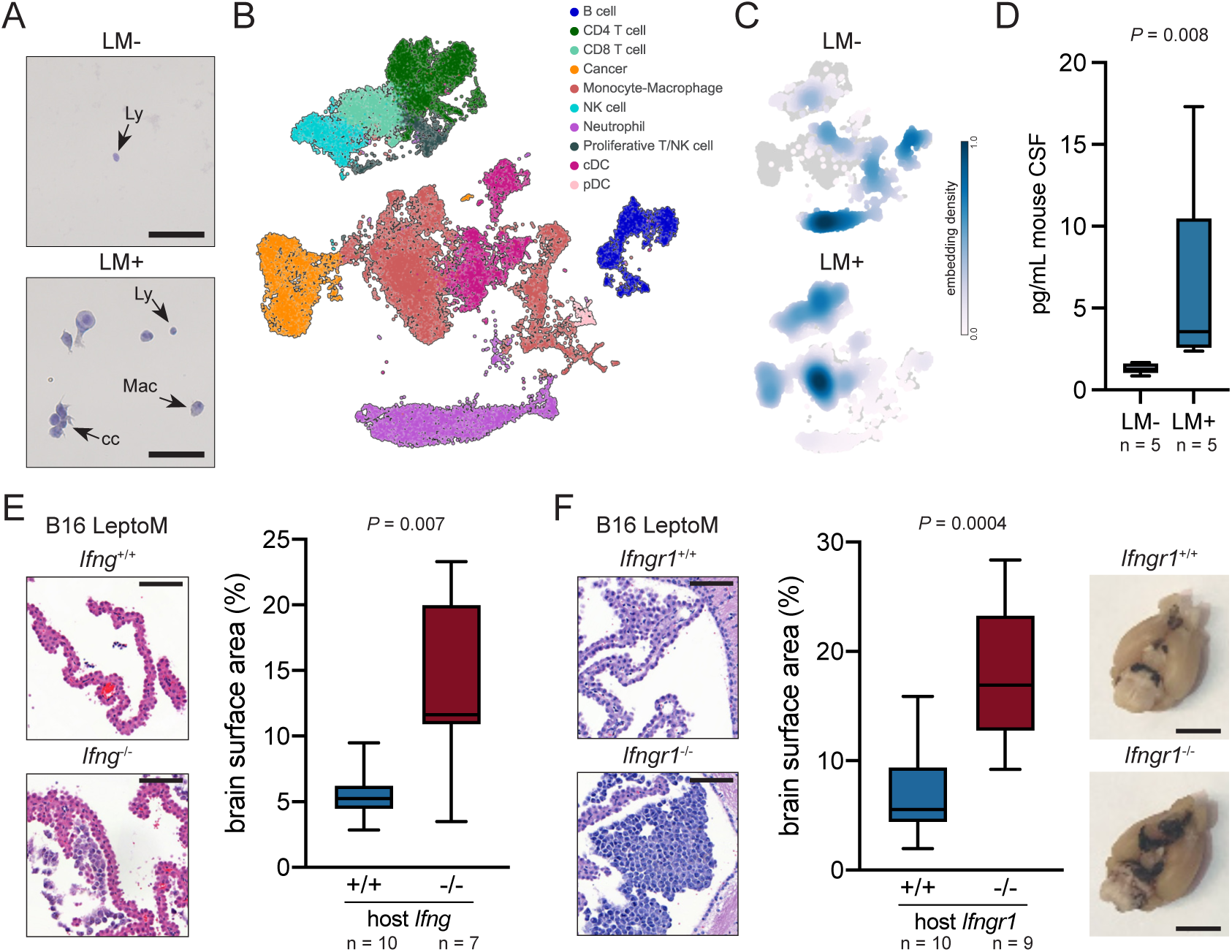
Host IFN-γ signaling suppresses expansion of immunocompetent mouse LM cells. (**A**) Representative images of hematoxylin-stained cytospins from vehicle-(top) and E0771 LeptoM-injected mice (bottom) with major cell populations indicated as Ly - lymphocyte, Mac - monocyte-macrophage, cc - cancer cell (n = 3 *per* group, scale bar = 50 μm). See fig. S3 for characterization of immunocompetent LeptoM mouse lines. (**B**) UMAP of cellular material isolated from vehicle- and LLC LeptoM-injected mice two weeks after inoculation, subjected to single-cell proteogenomic profiling with 10x CITE-seq (n = 7,528 cells from vehicle-injected and n = 19,534 cells from LLC LeptoM-injected mice, n = 6 animals *per* group). Cell type annotations are provided in fig. S4, and experiment overview is provided in fig. S15. (**C**) Embedding density plots from LM- and LM+ mice, showing relative cell type abundance *per* condition, projected onto UMAP. (**D**) Levels of IFN-**γ** in the CSF collected from naïve or LeptoM-bearing mice, detected by cytometric bead array. (**E**) Representative leptomeningeal tissue sections stained with H&E (scale bar = 100 μm). Box plot illustrates brain surface area covered with pigmented B16 LeptoM cells delivered intracisternally into C57BL/6 *Ifng*-proficient and -deficient animals, quantified two weeks after injection. (**F**) Representative leptomeningeal tissue sections stained with H&E (scale bar = 100 μm). Box plot illustrates brain surface area covered with pigmented B16 LeptoM cells delivered intracisternally into C57BL/6 *Ifngr1*-proficient and -deficient animals, quantified two weeks after injection. Photographs show the involvement of mouse basilar meninges with plaques of B16 LeptoM melanoma cells (scale bar = 5 mm).

In light of this myeloid predominance, we investigated leptomeningeal IFN-**γ** signaling ^11^. We detected elevated levels of IFN-**γ** in mouse LM, compared to vehicle-injected animals (**Fig. 2D**), analogous to human disease (**Fig. 1D**). To identify its source, we subjected CSF collected from mice with and without LM to flow cytometric assessment of IFN-**γ** production (**fig. S5A**). We find that leptomeningeal T and NK cells produce IFN-**γ**. In parallel, we queried our human and mouse single-cell atlases for *IFNG* transcript. In both mouse and human leptomeninges, T cells and NK cells produce IFN-**γ** (**fig. S5B**). Because IFN-**γ** binding to its cognate receptors triggers a signaling cascade that results in phosphorylation of STAT1 (pSTAT1) ^12^, we assessed the levels of pSTAT1 in leptomeningeal immune infiltrates by flow cytometry. We detected increased levels of pSTAT1 in mouse leptomeningeal dendritic cells, monocyte-macrophages, and T cells, but not natural killer (NK) cells (**fig. S5C**). Taken together, these results support a model whereby leptomeningeal monocyte-macrophages, dendritic cells and T cells respond to IFN-**γ** generated by leptomeningeal T and NK cells. To assess the contributions of IFN-**γ** signaling to leptomeningeal cancer growth, we leveraged transgenic host mice lacking either the sole type II Interferon ligand, *Ifng*, or its receptor, *Ifngr1*, resulting in whole-body impairment of IFN-**γ** signaling ^13, 14^. In both transgenic hosts and the three tested LM models, interruption of IFN-**γ** signaling led to uncontrolled cancer cell growth in the leptomeninges (**Fig. 2, E and F, and fig. S6**). This effect was not observed when these LeptoM cells were orthotopically implanted in their primary sites or the subcutaneous tissues (**fig. S7**), consistent with a leptomeningeal-specific role for IFN-**γ**.

In a context-dependent fashion, IFN-**γ** may either promote or inhibit cancer growth. This can be the result of direct IFN-**γ** signal to the cancer cell, or indirect signaling to the tumor microenvironment. To investigate whether IFN-**γ** acts directly on cancer cells and supresses their growth *in vivo*, we next genetically abrogated IFN-**γ** signaling in cancer cells by knocking out the *Ifngr2* subunit of IFN-**γ** receptor with CRISPR/Cas9. Unlike control clones, these knock-out lines were unable to propagate IFN-**γ** response that normally leads to upregulation of MHC class I on the cell surface (**fig. S8, A to C**). The lack of *Ifngr2* in these cells did not alter their growth *in vitro* (**fig. S8, D to F**), or *in vivo* (**fig. S8, G to I**). Cancer-intrinsic IFN-**γ** signaling is thus not required for cancer cell survival in the leptomeninges. Therefore, IFN-**γ** mediates leptomeningeal cancer cell growth through indirect effects on the microenvironment. Because knockout of host IFN-**γ** promoted cancer cell growth, we pursued a complementary add-back strategy with weekly intra-cisternal introduction of recombinant mouse IFN-**γ**. While LeptoM cancer cells demonstrate capacity to receive receive IFN-**γ** signals (**fig. S9, A to D),** this does not significantly impact their proliferation *in vitro* (**fig. S9, D to F**). However, *in vivo*, addition of IFN-**γ** suppressed cancer cell growth within the leptomeninges (**fig. S9, F to J**). Thus, IFN-**γ** suppressed intrathecal cancer cell growth in an indirect fashion, suggesting an interplay between IFN-**γ** and other cells in this inflammatory microenvironment.

### Leptomeningeal interferon-**γ** tumor suppression is independent of antigen presentation

To uncover the downstream IFN-**γ** effectors in the context of LM, we designed an experimental system enabling manipulation of CSF composition without frequent anesthesia or injection of foreign agents into the leptomeninges. We constructed an adeno-associated viral (AAV)-based expression system to induce expression of exogenous *Ifng* or a control gene, *Egfp*, specifically in the mouse leptomeninges ^15, 16^, (**fig. S10, A to B**). With this technique, overexpressed leptomeningeal IFN-**γ** resulted in dramatic control of leptomeningeal metastatic cancer cell growth in all six syngeneic LeptoM models; overexpressed EGFP did not (**Fig 3, A to F, and fig. S10, C to H**). Importantly, this overexpression system did not result in neurodegeneration or neuroinflammation, as in the case of Type I Interferons ^17^. Indeed, we observed a normal profile of astrocytes lining the ventricular space, without apparent activation of parenchymal microglia, depletion of neural progenitors, change in neuronal tract distribution, or change in mature cortical neuron numbers (**fig. S11**). Similar to earlier reports, we detected a decrease in the immature oligodendrocyte population in *corpus callosum* ^18, 19^. This was not reflected in cortical and subcortical layers where we detected only a minor decrease in differentiated, CNPase-positive cortical and subcortical oligodendrocytes (**fig. S12**).

**Fig. 3.**
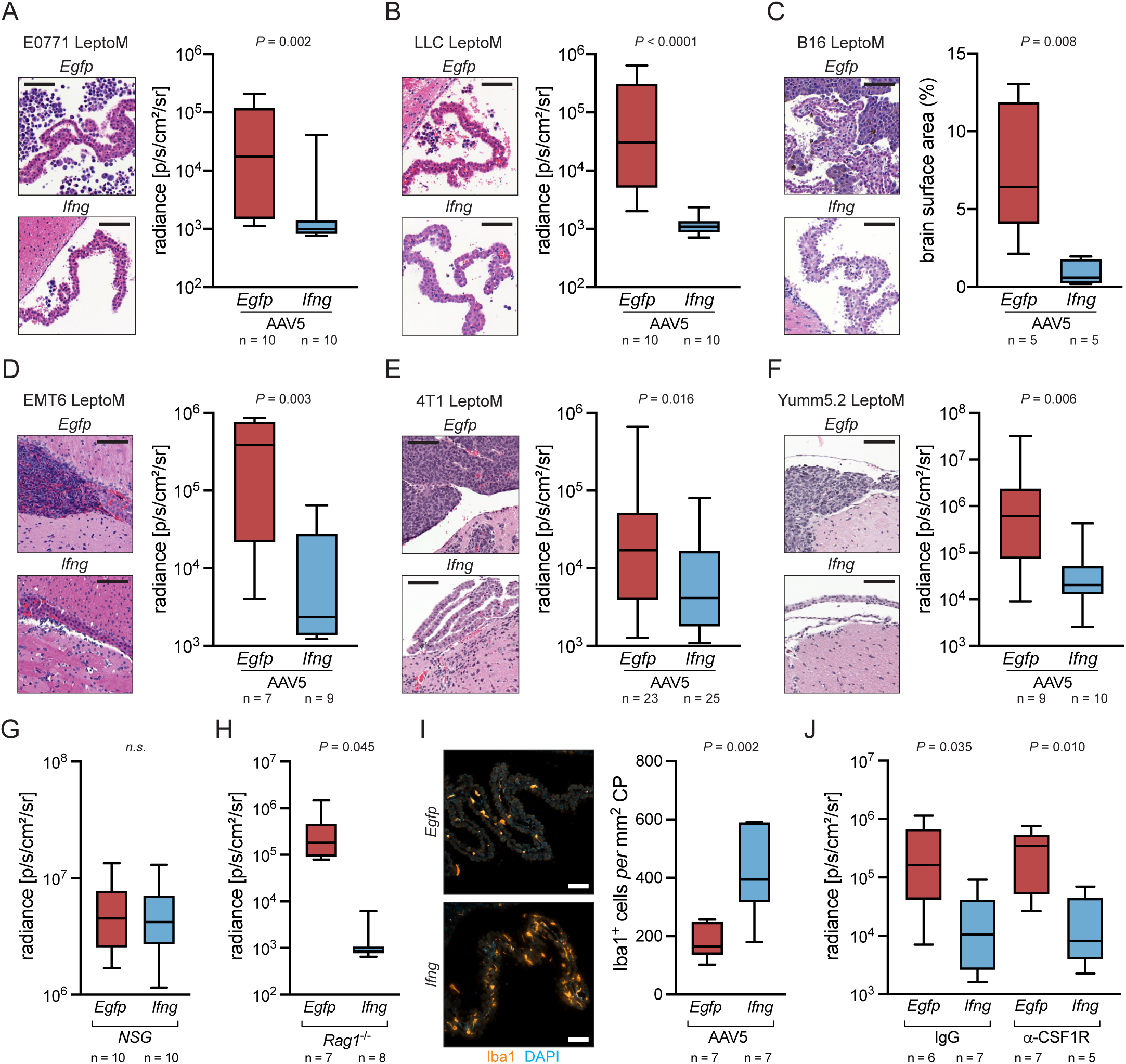
IFN-γ controls the growth of metastatic cancer in leptomeninges independent of the adaptive immune system and monocyte-macrophages. (**A**) Representative leptomeningeal tissue sections stained with H&E (scale bar = 100 μm). Box plot illustrates *in vivo* radiance of E0771 LeptoM cells delivered intracisternally into C57Bl/6-*Tyr*^c-2^ animals overexpressing *Egfp* or *Ifng* in the leptomeninges, quantified two weeks after injection. (**B**) Representative leptomeningeal tissue sections stained with H&E (scale bar = 100 μm). Box plot illustrates *in vivo* radiance of LLC LeptoM cells delivered intracisternally into C57Bl/6-*Tyr*^c-2^ animals overexpressing *Egfp* or *Ifng* in the leptomeninges, quantified two weeks after injection. (**C**) Representative leptomeningeal tissue sections stained with H&E (scale bar = 100 μm). Box plot illustrates brain surface area covered with pigmented B16 LeptoM cells delivered intracisternally into C57BL/6 animals overexpressing *Egfp* or *Ifng* in the leptomeninges, quantified two weeks after injection. (**D**) Representative leptomeningeal tissue sections stained with H&E (scale bar = 100 μm). Box plot illustrates *in vivo* radiance of EMT6 LeptoM cells delivered intracisternally into BALB/c animals overexpressing *Egfp* or *Ifng* in the leptomeninges, quantified two weeks after injection. (**E**) Representative leptomeningeal tissue sections stained with H&E (scale bar = 100 μm). Box plot illustrates *in vivo* radiance of 4T1 LeptoM cells delivered intracisternally into BALB/c animals overexpressing *Egfp* or *Ifng* in the leptomeninges, quantified one week after injection. (**F**) Representative leptomeningeal tissue sections stained with H&E (scale bar = 100 μm). Box plot illustrates *in vivo* radiance of Yumm5.2 LeptoM cells delivered intracisternally into C57Bl6-*Tyr*^c-2^ animals overexpressing *Egfp* or *Ifng* in the leptomeninges, quantified three weeks after injection. (**G**) *In vivo* radiance of LLC LeptoM cells delivered intracisternally into NSG animals overexpressing *Egfp* or *Ifng* in the leptomeninges, quantified two weeks after injection. (NSG - non-obese, diabetic, severe combined immunodeficient, *Il2rg*^null^). (**H**) I*n vivo* radiance of LLC LeptoM cells delivered intracisternally into Rag1-deficient animals overexpressing *Egfp* or *Ifng* in the leptomeninges, quantified two weeks after injection. (NSG - non-obese, diabetic, severe combined immunodeficient, *Il2rg*^null^). (**I**) Representative immunofluorescent image of brain tissue from cancer-naïve animals overexpressing *Egfp* or *Ifng* in the leptomeninges, stained for Iba1^+^ myeloid cells (scale bar = 50 μm). Box plot illustrates quantification of Iba1^+^ cells in the ventricular choroid plexi. (**J**) I*n vivo* radiance of LLC LeptoM cells delivered intracisternally into C57Bl6-*Tyr*^c-2^ animals overexpressing *Egfp* or *Ifng* in the leptomeninges and tri-weekly infused with non-targeting isotype control antibody or CSF1R-targeting antibody, quantified two weeks after injection.

With this tool in hand, we set to identify the key cell population(s) responsible for IFN-**γ** - dependent cancer control in the leptomeninges. The anti-cancer effect of IFN-**γ** was diminished when this IFN-**γ** overexpression system was established in fully immunodeficient NSG mice, confirming that immune cells mediate IFN-**γ**’s anti-cancer activity in the leptomeninges (**Fig. 3G, and fig. S13, A to C**). IFN-**γ** positively regulates antigen presentation ^12^. It was therefore surprising to observe the IFN-**γ** anti-tumor effect was preserved in *Rag1*-deficient animals with impaired adaptive immune system, indicating that IFN-**γ**’s anti-tumor function is independent of antigen presentation in the leptomeninges (**Fig. 3H, and fig. S13, D to F**). Iba1+ monocytes and macrophages are well-known IFN-**γ** effectors ^12^. Overexpression of *Ifng* resulted in accumulation of Iba1+ myeloid cells in the choroid plexus (**Fig. 3I**), a structure that acts as an interface between the periphery and the leptomeninges, produces the majority of CSF, and serves as a gateway for immune cell entry ^20, 21^. In our system, neither antibody-based, nor chemical depletion of monocyte-macrophage population resulted in impaired tumor growth control by IFN-**γ** (**Fig. 3J, and fig. S13, G and H**). IFN-**γ** -mediated leptomeningeal tumor control is thus dependent on the immune system, but independent of an antigen presentation, adaptive immunity, and monocyte-macrophage function. We therefore turned our attention to leptomeningeal dendritic cells.

### Dendritic cells orchestrate innate anti-tumor immune response in the leptomeninges

Conventional DCs (cDC) are a professional phagocytic myeloid immune cell lineage that can propagate IFN-**γ** response. Their function in an antigen-independent setting is, however, less explored. To specifically deplete the cDC lineage in the mouse and clarify their role in LM, we took advantage of a transgenic line that expresses human diphtheria toxin receptor (DTR) under the control of endogenous mouse *Zbtb46* ^22^. Within the hematopoietic compartment the ZBTB46 expression is restricted specifically to cDC progenitors, it is also expressed by other body cell types, such as endothelium. To avoid consequences related to systemic depletion of ZBTB46-expressing cells, we generated bone marrow chimeras: We infused lethally irradiated wild-type recipient mice with bone marrow from wild-type or ZBTB46-DTR animals. In this scenario, diphtheria toxin (DTx) eliminates ZBTB46- expressing cDC progenitors (**fig. S14**), while retaining the normal function of other, non-hematopoietic cell types. After reconstitution of normal bone marrow function, we overexpressed leptomeningeal *Ifng* or *Egfp*, and introduced cancer cells. Introduction of DTx into wild-type chimera hosts did not alter the activity of IFN-**γ**; mice with ablated cDC demonstrated reduced IFN-**γ** -dependent tumor control (**Fig. 4A**). These experiments suggested that leptomeningeal DCs, responding to IFN-**γ** (**fig. S5C**), mediate its anti-tumor action.

**Fig. 4.**
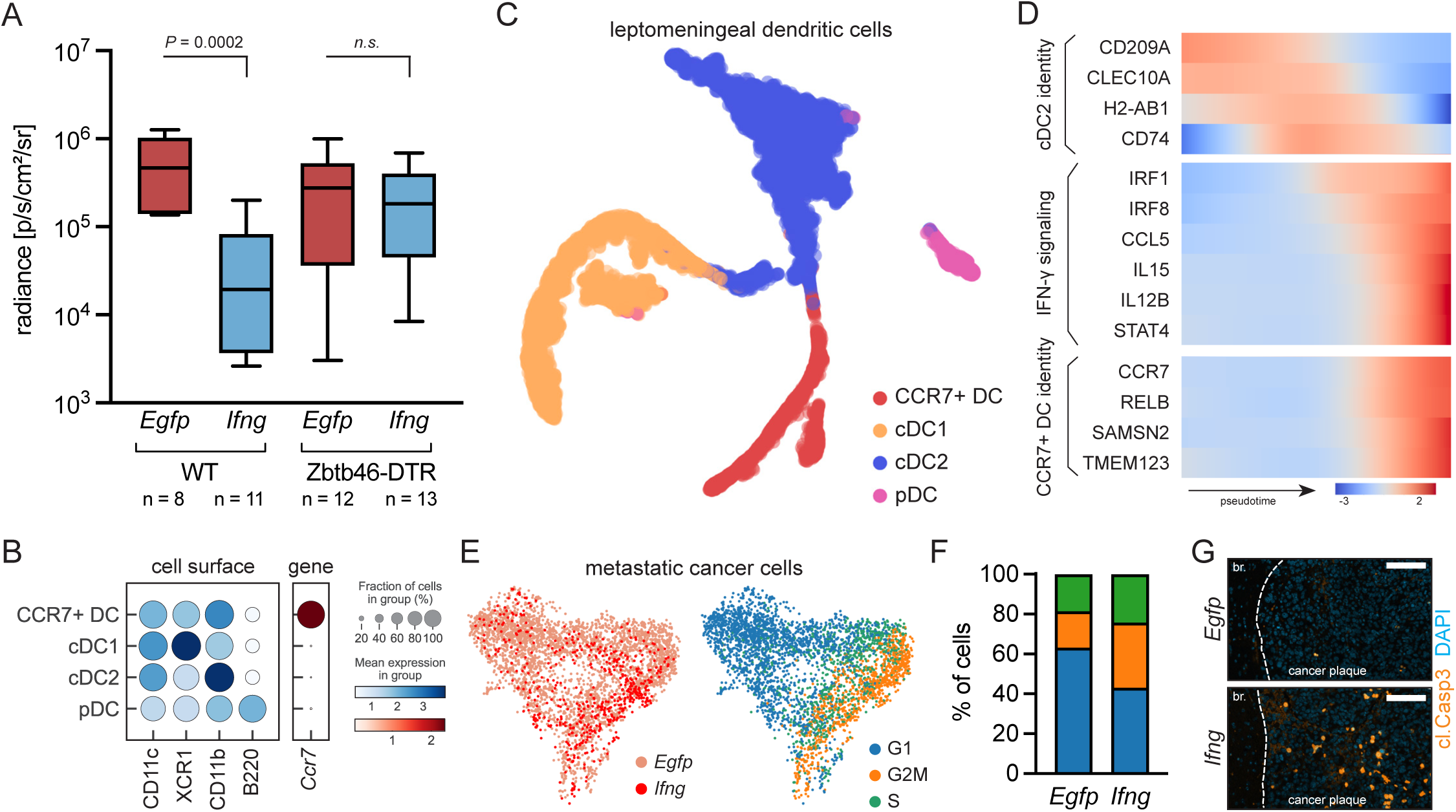
Leptomeningeal IFN-γ supports cDC maturation. (**A**) *In vivo* radiance of LLC LeptoM cells delivered intracisternally into C57Bl6-*Tyr*^c-2^ bone marrow chimeras overexpressing *Egfp* or *Ifng* in the leptomeninges after administration of diphteria toxin. Host mice were infused with bone marrow from wild-type or *Zbtb46-*DTR^+/+^ C57Bl6 mice. Depletion efficiency is quantified in fig. S14. (**B**) Dot plot showing expression of characteristic dendritic cell (DC) surface proteins and *Ccr7* gene, as determined with single-cell proteogenomics (dendritic cells pooled from 4 conditions, and 6 mice *per* condition were included; see fig. S15-S17 for details). (**C**) tSNE projection of mouse leptomeningeal DC types with predicted maturation streamlines from vehicle- and LLC LeptoM-injected mice two weeks after inoculation, subjected to 10x CITE-seq (total of n = 7,566 cells pooled from 4 conditions and n = 6 animals *per* group). cDC and pDC cells shown in fig. S15 were subsetted and reclustered, and tSNE was built in multiscale space derived from diffusion components, see Methods. (**D**) Expression trends in genes associated with cDC2 identity, IFN-**γ** signaling, and CCR7+ identity, along diffusion pseudotime axis representing cDC2-CCR7+ DC maturation. Gene trends were computed with Palantir ^27^. See also fig. S16 and S17, and Methods. (**E**) UMAP projection of identity (left) and cell cycle phase prediction (right) in LLC LeptoM cancer cells, isolated from animals overexpressing *Egfp* or *Ifng* in the leptomeninges (n = 3,161 and n = 557 cells from *Egfp* or *Ifng*-overexpressing mice, respectively; n = 6 animals *per* group). Cancer cells were identified as keratin- and CD63-expressing cluster and visualized without additional re-clustering, see fig. S4 and S15. (**F**) Quantification of LLC LeptoM gene expression-based cell cycle prediction from panel E. See also fig. S18, A to C. Predictions were computed using scores of gene lists characteristic for S and G2/M phases, see Methods. (**G**) Representative immunofluorescent image of E0771 LeptoM cancer plaques in animals overexpressing *Egfp* or *Ifng* in the leptomeninges, stained for cleaved Caspase 3 (scale bar = 50 μm). For quantification see fig. S18, D to F.

To capture the complexity of the IFN-**γ** response in the leptomeningeal space at a systems level, we isolated leptomeningeal cells from *Egfp-* and *Ifng-*overexpressing mice in the presence and absence of LM, and profiled these cells with CITE-seq (**fig. S15**; total n = 24 mice from 4 conditions). We confirmed the presence of all classical DC populations: cDCs1 and cDCs2, migratory CCR7+ DCs, and plasmacytoid DCs (pDCs; **Fig. 4B and C**). Molecular profiling of DCs isolated from *Egfp-* and *Ifng-*overexpressing mice revealed striking similarities between mouse and human leptomeningeal DCs (**fig. S16**), as well as site-specific (leptomeningeal) imprinted expression patterns different from those observed within extracranial sites ^23, 24^. In the presence of cancer, or after IFN-**γ** induction, cDC populations accumulate within the leptomeninges (**fig. S17, A to C**). To address IFN-**γ** - dependent relationships between these cDC populations, we queried our proteogenomic atlas. Outside of the CNS, CCR7+ DCs can arise from both cDC1 and cDC2 populations ^25^. The majority of leptomeningeal CCR7+ DCs, however, retained of the cDC2 surface expression profile, as detected with CITE-seq (**fig. S17, D and E**). Given the leptomeningeal-specific expression pattern, we elected to approach this computationally and first employed CellRank to predict terminal cell states, without the need to indicate the initial cell (**fig. S17F**) ^26^. This analysis identified cDC2 cells as the major contributors to the leptomeningeal CCR7+ DC pool; it also identified CCR7+ DCs as predominantly a product of cDC2 maturation (**fig. S17, G and H**). We then reproduced trajectory analyses with Palantir, modeling the cDC2-CCR7+ DC maturation axis (**fig. S17, I and J**) ^27^. We detected enrichment of IFN-**γ** -associated genes as cells transition to CCR7+ DCs, consistent with IFN-**γ** contribution to CCR7+DC maturation from cDC2 cells (**Fig. 4D**) ^28^. Because the anti-tumor effect of leptomeningeal IFN-**γ** does not rely on antigen presentation, we examined other anti-tumor pathways including cancer cell proliferation and death. Prediction of cell cycling in transcriptomic cancer cell data revealed that cancer cells isolated from *Ifng*-overexpressing mice did not show defective proliferation (**Fig. 4, E and F, and fig. S18, A to C**). However, immunofluorescence of cancer cells in the leptomeninges identified elevated caspase expression in the *Ifng*-overexpressing animals, consistent with apoptotic cell death (**Fig. 4G** **and fig. S18, D to F**). These results suggested that a cytotoxic immune population, supported by cDCs, restricts cancer cell expansion in the leptomeninges.

### Dendritic cell-generated cytokines drive proliferation of leptomeningeal NK cells

We therefore turned our attention to the transcriptomic profiles of leptomeningeal NK cells, cytotoxic effectors capable of tumor cell killing ^29^. Mouse leptomeninges contained naïve, activated, and proliferating NK cells. In the presence of cancer, a minor population of senescent NK cells was also apparent (**Fig. 5, A and B, and fig. S19, A to C**). Human CSF demonstrated analogous populations of naïve-like and activated-like NK cells (**fig. S19, D to G**). Independent of cancer, leptomeningeal *Ifng* overexpression induced increased NK cell proliferation; this effect was retained in NK cells isolated from *Ifng* overexpressing cancer-bearing animals (**Fig. 5C**).

**Fig. 5.**
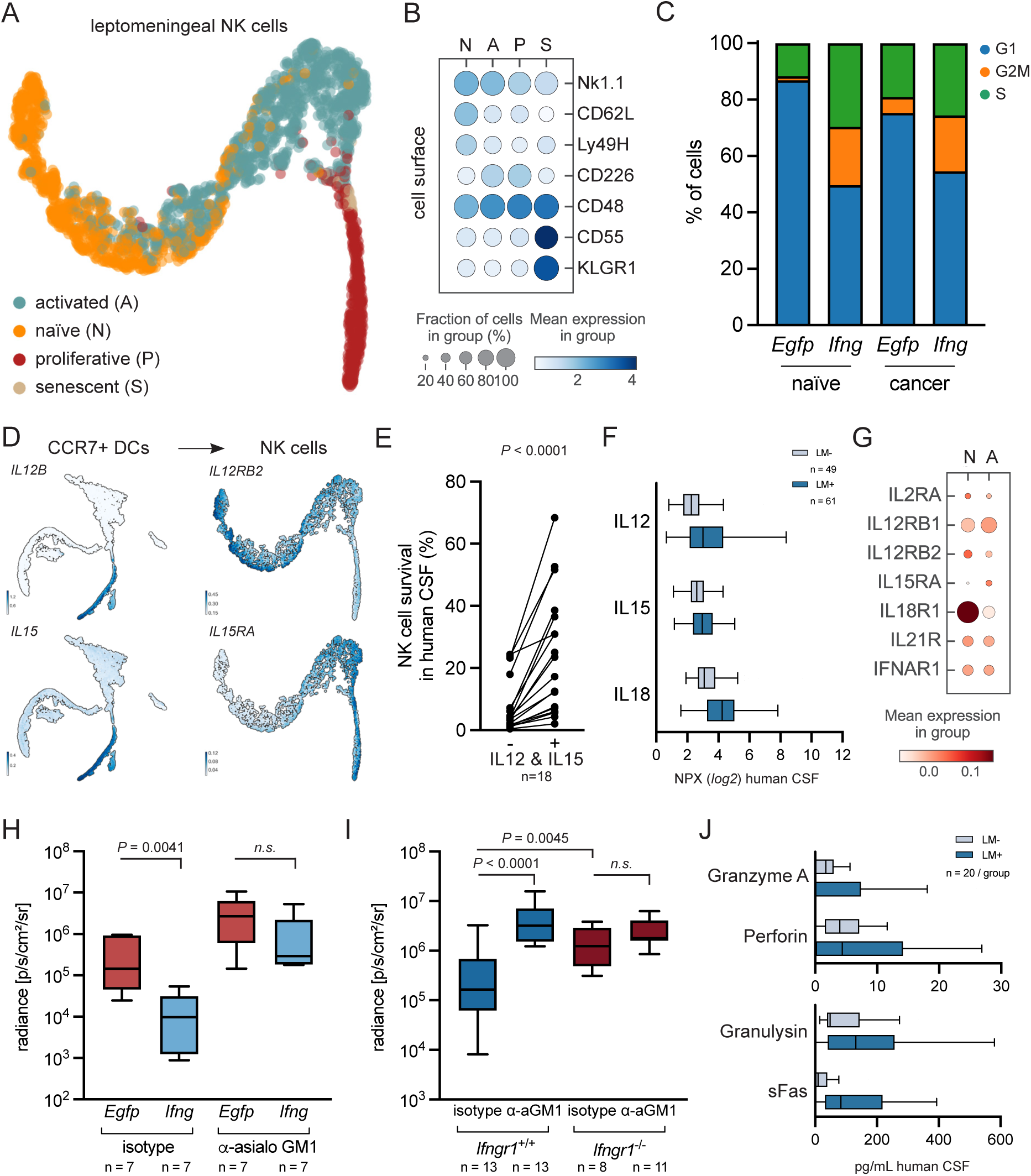
cDC-derived cytokines mediate NK cell activity and proliferation to prevent cancer cell outgrowth. (**A**) tSNE projection of mouse leptomeningeal natural killer (NK) cell states from vehicle- and LLC LeptoM-injected mice two weeks after inoculation, subjected to 10x CITE-seq (total of n = 2,247 cells from 4 conditions, n = 6 animals *per* group). Nk1.1^+^ *NKG7*^+^ CD3^-^ TCRý^-^ cells from fig. S15 were subsetted and reclustered, and tSNE was built in multiscale space derived from diffusion components, see Methods. (**B**) Expression of cell state-enriched NK surface proteins in mouse, as determined with single-cell proteogenomics. (**C**) NK cell cycle prediction in vehicle-injected, cancer-naïve or LLC LeptoM-bearing animals overexpressing *Egfp* or *Ifng* in the leptomeninges, as determined with single-cell proteogenomics. Predictions were computed using scores of gene lists characteristic for S and G2/M phases, see Methods. (**D**) Smoothened gene expression of selected CCR7+ DC ligands and NK cell receptors, projected onto tSNE plots. Gene imputation was performed with Markov affinity-based graph imputation of cells (MAGIC) ^43^. (**E**) Paired analysis of NK cell survival in human LM- CSF without or with addition of recombinant mouse IL12 and IL15 (results pooled from four independent replicates, paired *t* test). (**F**) Relative abundance of IL12, IL15, and IL18 in the CSF of LM- and LM+ cancer patients, as determined with targeted proteomics. (**G**) Expression of cell state-enriched human NK surface proteins, as determined with single- cell transcriptomics. For annotation see fig. S19, D to F. (**H**) *In vivo* radiance of LLC LeptoM cells delivered intracisternally into C57Bl6-*Tyr*^c-2^ animals overexpressing *Egfp* or *Ifng* in the leptomeninges and bi-weekly infused with non-targeting isotype control antibody or asialo-GM1-targeting antibody, quantified two weeks after injection. (**I**) *In vivo* radiance of LLC LeptoM cells delivered intracisternally into C57BL/6 *Ifngr1*- proficient and -deficient animals and bi-weekly infused with non-targeting isotype control antibody or asialo-GM1-targeting antibody, quantified two weeks after injection. (**J**) Quantification of soluble Granzyme A, Perforin, Granulysin, and sFas in the CSF of LM- and LM+ cancer patients, as determined by cytokine bead arrays.

We next examined communication between leptomeningeal CCR7+ DCs and NK cells. As determined by CITE-seq, mouse leptomeningeal CCR7+ DCs specifically produced IL12 and IL15, two cytokines that promote survival and proliferation of NK cells; leptomeningeal NK cells expressed their cognate receptors (**Fig. 5D**). To examine this putative cell-cell communication, we cultured mouse splenic NK cells in human CSF isolated from patients without LM. CSF represents a notoriously nutrient-sparse environment with minimal growth factors ^30^. Within CSF, naïve splenic NK cell survival was impaired; this effect was rescued by the addition of recombinant mouse IL12 and IL15 (**Fig. 5E**). Mirroring findings in our mouse models, we detected increased levels of NK cell-supporting cytokines in the CSF from patients harboring LM (**Fig. 5F**), as well as transcripts of their receptors in human leptomeningeal NK cells (**Fig. 5G**). To demonstrate the role of NK cells in IFN-**γ** -dependent cancer control, we depleted NK cells in mice overexpressing *Ifng* in our AAV5 system (**fig. S10, A and B**). As expected, we observed control of tumor growth and extended survival in mice treated with control antibody in the presence of leptomeningeal IFN-**γ** (**Fig. 5H, and fig. S20**). This phenotype was abolished in mice with antibody-depleted NK cells, supporting a model whereby NK cells serve as the leptomeningeal effector cells in the context of IFN-**γ**. We next depleted NK cells in *Ifngr1^-/-^* host mice. In this epistasis experiment, NK cell depletion in mice with non-functional IFN-**γ** signaling did not further accelerate leptomeningeal cancer cell growth, confirming that IFN-**γ** signaling precedes NK cell-dependent cancer elimination (**Fig. 5I**). We uncovered evidence of NK cell activation in human LM in the form of elevated levels of granzyme A, perforin, granulysin, and sFas as well as enrichment of activated NK cells in the CSF of LM patients (**Fig. 5J** **and fig. S19, D and F**). Taken together, our data are consistent with a model whereby NK cell- and T cell-derived leptomeningeal IFN-**γ** acts on cDCs, supporting their maturation into CCR7+ DCs. These cells then produce a spectrum of lymphocyte-supporting cytokines, promoting NK cell proliferation and anti-leptomeningeal tumor action (**Fig. 6**).

**Fig. 6.**
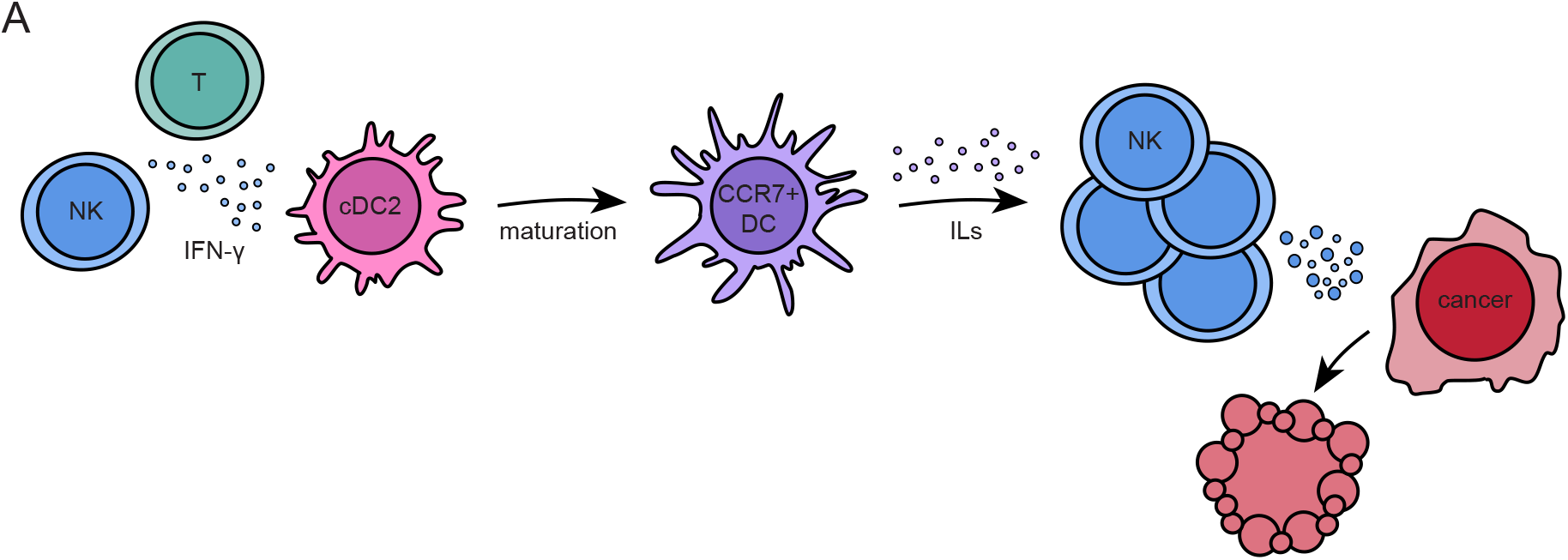
Leptomeningeal dendritic cells represent the essential IFN-γ target. (**A**) Schematic highlighting the main findings of this study. Leptomeningeal IFN-**γ**, produced mainly by T and NK cells, supports maturation of conventional DC2 into migratory DCs. These migratory DCs are characterized by the expression of *Ccr7* in mouse, and *LAMP3* in human. In an antigen-independent manner, these newly raised leptomeningeal migratory DCs produce an array of interleukins that support survival and proliferation of NK cells. NK are the cytotoxic effectors that control the expansion of metastatic cells in leptomeninges.

## Discussion

We have defined the molecular interactions between metastatic cancer and immune cells within the leptomeninges. To capture this complex oncologic ecosystem, we have employed single cell transcriptional and proteomic profiling of clinical samples. In doing so, we identified IFN-**γ** as a key mediator of anti-cancer response within the leptomeninges. To mechanistically dissect the growth suppressive action of leptomeningeal IFN-**γ**, we generated several new immunocompetent animal models of LM. We found that although leptomeningeal IFN-**γ** attracts myeloid cells into the leptomeningeal space, it does not promote anti-tumor activity in the macrophage population. Rather, leptomeningeal IFN-**γ** targets dendritic cells, promoting cDC2 maturation. Surprisingly, these dendritic cells orchestrate anti-cancer activity in an antigen-independent manner, generating cytokine signals to support the cytotoxic action of natural killer cells.

LM represents a fundamentally inflammatory pathology. Indeed, LM was originally described as a “carcinomatous meningitis”, reflecting the characteristic abundant immune infiltrate and the purulent exudate found at autopsy ^31, 32^. Recent work demonstrates that certain aspects of leptomeningeal inflammation can support cancer cell growth: Cancer cells within the leptomeningeal space respond to IL-8 and IL-6 to transcribe the iron binding and iron transport genes LCN2 and SLC22A17 ^4^; Cancer cell-generated complement C3 disrupts the blood-CSF-barrier to enrich the CSF and support cancer cell growth in the space ^1^. However, inflammatory signaling in the leptomeninges does not universally support LM. In this report, we have uncovered leptomeningeal inflammatory signaling that can interrupt cancer cell growth: IFN-**γ**. We identified elevation of leptomeningeal IFN-**γ** as a hallmark of LM-induced pleocytosis across multiple tumor types. Moreover, higher CSF IFN-**γ** at diagnosis portends a more favorable prognosis for these patients.

IFN-**γ** is a classical tumor suppressive cytokine derived predominantly by Th1 CD4+ T cells, as well as CD8+ T cells, NK cells, NKT cells, and minor population of other immune cell types ^33^. Investigation of IFN-**γ** within the leptomeningeal space revealed anatomically distinct features: the proportion of immune cells expressing this protein, or its transcript, appeared to be insufficiently low even at in the absence of malignancy, suggesting that the leptomeninges actively maintain low production of this pleiotropic cytokine, possibly to impede neurotoxicity interferon ligands ^7, 17^. IFN-**γ** stimulates the recruitment of a wide variety of immune cell types into the tumor microenvironment, particularly through the upregulation of CXC chemokines CXCL9, -10, and -11 ^33^. The impressive pleocytosis in LM patients and experimental animal models can be, to some extent, explained by accumulation of these IFN-**γ** -regulated chemokines. However, both CC and CXC chemokines are dramatically elevated in the leptomeninges of patients harboring systemic inflammation or prolonged COVID-19, yet their accumulation does not necessarily result in clinically relevant CSF pleocytosis, suggesting additional level of immune cell entry control into the CSF ^2, 4, 7, 34^. Why CSF IFN-**γ** and its downstream ligands do not consistently result in leptomeningeal accumulation of immune cells remains an open question.

In immunocompetent settings, IFN-**γ** prevents the establishment of spontaneous and chemically-induced tumors by enhancing cancer cell recognition, increasing antigen processing via MHC class I and II in the extracranial sites and brain ^35–37^. Unlike other anatomic compartments, the tumor-suppressive role of IFN-**γ** within the leptomeninges was unexpectedly independent of both the adaptive immune system and monocyte-macrophages. Instead, leptomeningeal DCs represent the essential IFN-**γ** target. Our systems-level approach suggests that metastasis renders the leptomeningeal space an unusually dendritic cell-rich environment, when compared to extracranial sites with relatively sparse proportion of infiltrating dendritic cells. Indeed, cytometric analysis of STAT1 phosphorylation in the presence of LM was most apparent in DCs. Moreover, single-cell proteogenomic analysis of leptomeningeal DCs further suggested a previously underappreciated role of IFN-**γ**: cDC maturation into CCR7+, migratory dendritic cells. Trajectory analysis of mouse cDCs support the assertion that these CCR7+ DCs are predominantly a product of cDC2 maturation. In other, extracranial tumors, both cDC1 and cDC2 equally contribute to the migratory DC pool ^25^. In antigen-independent settings, these migratory DCs produce an array of immune cell pro-survival and proliferation factors. In the harsh leptomeningeal environment, these DC-generated signals are necessary to sustain effector cell viability and activation. Indeed, we show that NK cells proliferate more in the setting of *Ifng* overexpression, and that this is supported by the presence of migratory DC-derived signals including IL12 and IL15.

Improved understanding of LM specific cancer cell-immune cell interactions suggests novel approaches to immune-oncology within the CNS and prompts a more nuanced view of the immune system in the leptomeningeal space. Our findings demonstrate that leptomeningeal metastatic cancer cell growth is largely controlled by the innate immune system. This may explain disappointing outcomes in LM-focused clinical trials targeting adaptive immunity ^38–40^. We propose that DC and NK cell-engaging therapies - both already showing promising clinical results in solid tumors and especially hematologic malignancies - may act as more rational strategies to control of this bleak complication of cancer ^41, 42^.

## Material and data availability

RNA-seq datasets were deposited online in the NCBI Gene Expression Omnibus (GEO) under the accession numbers GSE221358 (bulk RNA-seq), GSE221593 (mouse single-cell proteogenomics), GSE221522 (human single-cell RNA-seq). Commercially available materials can be obtained from vendors. Materials generated in this study are available from the corresponding author upon signing the MSKCC Material Transfer Agreement. Human samples used in this study are limited biological resource, not available for further distribution.

## Acknowledgements.

We wish to express our deep gratitude to the patients and families that inspire this work and donate clinical samples for this research. We also thank Anna Skakodub, Rachel Estrera, and Isaiah Osei-Gyening for help with management of human CSF bank, Dr. Chrysothemis Brown, Dr. Cassandra Burdziak, Dr. Lisa DeAngelis, Dr. Morgan Freret, Dr. Andrea Schietinger, Roshan Sharma, Dr. Thomas Walle, and Dr. Jedd Wolchok for fruitful discussions, to Dr. Majdi Alghader, Camille Derderian, and David Guber for technical assistance, and to Ojasvi Chaudhary and Ignas Masilionas for help with wet lab single-cell processing.

Financial support for this research was provided to A.B. by the Damon Runyon Cancer Research Foundation, the Pershing Square Sohn Cancer Research Alliance, the Alan and Sandra Gerry Metastasis and Tumor Ecosystems Center, and the NCI R01CA245499. J.R. was supported by the American Brain Tumor Association Basic Research Fellowship, the Terri Brodeur Breast Cancer Foundation, and the Fiona and Stanley Druckenmiller Center for Lung Cancer Research. All investigations were supported by NCI P30 CA008748 (Cancer Center Support Grant). This publication is part of the HTAN (Human Tumor Atlas Network) Consortium paper package. A list of HTAN members is available at https://humantumoratlas.org/htan-authors/.

## Author contributions

J.R. and A.B. conceived and designed the study, acquired funding, and wrote the manuscript. J.R. generated metastatic cell lines, developed the methodology, performed the experiments, analysed, and curated the data. X.T., M.L., D.I., and J.S. assisted with experiments and replicated critical experiments. J.R. and R.K. performed computational analysis under D.P.’s supervision. A.O. assisted with immunofluorescent staining and image analysis. K.C., J.A.W., and U.S. assisted with clinical annotations and human CSF collection. T.B. reviewed cytospin staining. R.C. supervised single-cell sequencing. All authors approved the final manuscript. A.B. supervised the study.

## Supplementary Materials and Methods

### Human CSF

Cancer patients undergoing routine clinical procedures including spinal tap, Ommaya reservoir tap, or a ventricular shunt provided informed consent. CSF collected in excess of that needed for clinical care was reserved for this use under MSKCC Institutional Review Board-approved protocols 20-117, 18-505, 13-039, 12-245, and 06-107. Human CSF was processed as described in ^1^, de-identified, and aliquoted. Cell-free CSF and CSF cell pellets were biobanked and stored at -80°C until further analysis. Patient medical records and MRI scans were reviewed to confirm the LM status by neurooncologists (U.S., J.A.W., and A.B.), and clinical data necessary for this study was abstracted and de-identified. Giemsa-stained cytospins were part of routine diagnostic assessment and were retrieved and reviewed by neuropathologist (T.B.).

### Human single-cell transcriptomics

#### Sample processing

Freshly collected CSF obtained by lumbar puncture was placed on ice and processed within two hours, as described previously ^1^, PBS-washed cells were encapsulated with Chromium Single Cell 3’ Library and Gel Bead Kit V2 (10x Genomics) and sequenced on an NovaSeq 6000 system (Illumina). Raw and pre-processed data were deposited to NCBI GEO under accession number GSE221522.

#### Data preprocessing, initial processing, and batch correction

Raw FASTQC files were pre-processed with SEQC ^2, 3^ with human reference genome hg38, and dense SEQC matrices were imported into Python. Each sample was plotted as a histogram of total counts *per* cell barcode on the *log* scale, resulting in a distribution with multiple modes and the threshold to remove the smallest mode, containing empty droplets and low-quality cells, was defined manually. We next removed any genes that had counts equal to 0 after filtering. To remove doublets, we run the DoubletDetection method (parameters n_iter = 50, p_thres = 1e-7, voter_thres = 0.8) ^4^. We outer joined the individual samples to keep all detected genes, filtered cells to a minimum count of UMI = 100 and minimum total expressed genes of 100. We initially detected 22,051 cells and , genes and retained 20,676 high-quality cells and 18,322 genes after filtering. We detected ∼1,497±898 genes per cell, ∼6,268±6,553 gene counts per cells, out of which 3.25±2.99% were mitochondrial genes (values represent mean ± one standard deviation). We normalized the library size, keeping raw count matrix for downstream analyses and removed any genes expressed in fewer than 5 cells. For downstream analysis, we further removed mitochondrial genes (prefix MT-), ribosomal genes (prefix RPS- or RPL-), and hemoglobin genes (prefix HB-). We run Scanorama (default settings; kNN = 20) ^5^ on the resulting AnnData object to batch correct across patients. Batch correction was validated as follows: (i) cancer cells have higher inter-patient heterogeneity (fig. S1), suggesting absence of overcorrection, and (ii) we identified and filtered out only few quasi-cancer cells from LM- patients after computational mixing (their presence was ruled out by pathologist during diagnostic cytology reading). This corrected matrix was employed for visualization, but not for individual gene comparisons. We then run PCA (sc.pp.pca, n_components = 100). We constructed a k-nearest neighbor graph (kNN) based on 30 nearest neighbors and 100 principal components, using the scanorama 100-dimensional matrix (instead of PCA matrix). We clustered the cells with Leiden (resolution 2.0) ^6^ and these Leiden clusters were merged according to major cell types, which were assigned based on marker gene expression, as showed in fig. S1. UMAP was computed with sc.tl.umap, using default parameters. The inter-patient heterogeneity was measured with Shannon entropy, *H_j_* (fig. S1, G to I) ^7^:

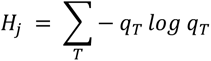

For each cell, the Shannon entropy measures the sample diversity of its nearest neighbors in the kNN graph. Each sample was subsampled to contain 500 cells. If samples are well-mixed, entropy of each cell will be high, while if samples are not well mixed entropies will tend to be low (this is true for cancer cells in general, which show extreme heterogeneity across patients). See fig. S1 for quality control plots. Human LM+ single-cell transcriptomic data was retrieved from NCBI GEO GSE150660 ^1^. Raw and pre-processed data are available through NCBI GEO under accession number GSE221522. All ten human samples were collected between December 2017 and May 2018 and processed with the same pipeline.

#### Subsetting of cells for downstream analyses and visualization

Subsetting was performed by selecting cell clusters from major Leiden populations, shown in Fig. 1B. For analysis of dendritic cells (DC), ‘cDC’ and ‘pDC’ clusters were subsetted and reclustered with sc.tl.umap and Leiden (resolution = 0.5). Cell type annotation was performed as follows: cDC1 cells are *CLEC9A*^+^*XCR1*^+^, cDC2 cells are *CLEC10A*^+^*CD1C*^+^, pDC cells are *IRF7*^+^*TCF4*^+^. Human LAMP3+ migratory dendritic cells are *LAMP3*^+^*CCR7*^+^ (orthologous to mouse CCR7+ DC). Two clusters bearing cDC2 signature were merged for further analyses. For analysis of natural killers (NK), ‘NK’ cluster was subsetted and reclustered with Leiden (resolution = 0.8), yielding in populations of cells with high *SELL* (CD62L) expression, further denoted as naïve-like, and populations with low *SELL* expression, denoted as activated-like and characterized by the expression of *CXCR6*. For analysis of both cell types, we run Palantir with default settings (n_components = 5, knn = 30) that allowed us to access MAGIC-imputed (Markov affinity-based graph imputation of cells) cell counts, and these imputed cell counts were used only for visualization with 2D plots ^8, 9^. UMAP, tSNE and heatmap plotting was performed using Scanpy ^10^ and scVelo ^11^ toolkits. Embedding density was computed with sc.tl.embedding_density (Fig. 1C). For NK cell gene expression heatmap, counts were first zero-centered with sc.pp.scale (fig. S19G). Code for pre-processing and downstream analysis is available from corresponding author and will be deposited to GitHub after peer review.

### Human CSF targeted proteomics

Samples were processed and analysed essentially as described in ^12^. Biobanked CSF collected between 2015-2020 was aliquoted and stored at -80°C at MSK Brain Tumor Center CSF Bank. Samples were slowly thawed on ice and 45 μL of CSF was mixed with 5 μL of 10% Triton X-100 (Sigma, T8787) in saline and incubated at room temperature for two hours (final concentration of Triton X-100 was 1%). Samples were then dispensed in a randomized fashion into 96-well PCR plates and stored at -80°C until further analysis. Relative levels of proteins in two targeted panels were detected using proximity extension assay (Olink Target 96 Inflammation and Olink Target 96 Neuro Exploratory, Olink). Additional control, LM- samples were retrieved from ^12^ (CoV- cohort). Protein abundance values are shown in NPX units (*log2* scale). The analytical range for each analyte is available online (www.olink.com).

### Mouse strains and housing

All animal studies were approved by the MSKCC Institutional Animal Care and Use Committee under the protocol 18-01-002. Wild-type C57Bl/6 (JAX#000664) were purchased from Jackson Laboratory or bred in-house. C57Bl/6-*Tyr*^c-2^ (JAX#000058, albino C57Bl/6) and BALB/c (JAX#000651) animals were purchased from the Jackson Laboratory. NSG animals were obtained from MSKCC RARC Colony Management Group. Purchased mice were allowed to habituate for at least one week before manipulation and experimentation. Transgenic lines on C57Bl/6 background were purchased from the Jackson Laboratory and bred in-house: *Ifng* knock-out line (B6.129S7-*Ifng^tm1Ts^*/J, JAX#002287), *Ifngr1* knock-out line (B6.129S7-*Ifngr1^tm1Agt^*/J, JAX#003288), *Rag1* knock-out line (B6.129S7-*Rag1^tm1Mom^*/J, JAX#002216), double-reported knock-in/knock-out *Cx3cr1*^GFP/GFP^*Ccr2*^RFP/RFP^ (B6.129(Cg)- *Cx3cr1^tm1Litt^ Ccr2^tm^*^2^*^.1Ifc^*/JernJ, JAX#032127). For homozygous breeding, breeding pairs and randomly selected progenies used in the experiments were genotyped as recommended. For experiments that involved bioluminescent imaging where wild-type animals were not compared to transgenic lines, albino C57Bl/6-*Tyr^c-2J^* animals were used. Mice in all experimental groups were age- (± 4 days), sex-, and fur color-matched. Mice used in this study were housed in a specific pathogen-free conditions, in an environment with controlled temperature and humidity, on 12-hour light/dark cycles (lights on/off at 6:00 am/pm), and with access to regular chow and sterilized tap water *ad libitum*.

### Cell culture

Mouse lung cancer LLC sublines were described previously ^13^. Mouse breast cancer E0771 cells were kind gift from Dr. Ekrem Emrah Er. B16-F10 (CRL-6475), Yumm5.2 (CRL-3367), EMT6 (CRL-2755), and 4T1 cells (CRL-2539) were obtained from ATCC. LentiX 293T cells (#632180) were obtained from Takara. PlasmoTest HEK Blue-2 cells (rep-pt1) were obtained from Invivogen. LLC, E0771, and B16 sublines and LentiX 293T and HEK Blue-2 cells were maintained in high-glucose DME (MSKCC Media Core), supplemented with 10% fetal bovine serum (FBS; Omega Scientific #FB-01) and 1% penicillin-streptomycin (P/S; Gibco #15140163) or 1x Primocin (Invivogen #ant-pm-2). Yumm5.2 sublines were maintained in high-glucose DME:F12 (MSKCC Media Core), supplemented with 10% FBS, 1% non-essential amino acids (Gibco #11140050) and 1% P/S or 1x Primocin. 4T1 sublines were maintained in RPMI (MSKCC Media Core), supplemented with 10% FBS and 1% P/S or 1x Primocin. EMT6 sublines maintained in Waymouth’s (MSKCC Media Core), supplemented with 10% FBS and 1% P/S or 1x Primocin. Cell lines were subcultured at least twice a week, replaced approximately after six weeks in culture with new stocks, stored in liquid nitrogen, and routinely tested negative for mycoplasma contamination. Proliferation of vehicle- or recombinant mouse IFN-**γ** -exposed (Biolegend #714006) cancer cells *in vitro* was measured with CellTiter-Glo luminescent cell viability assay (Promega G7572) 72 hours after seeding 500 cells *per* well into 96-well, white-walled plate (Corning).

### Genetic engineering of mouse cancer cell lines

Plasmid DNA was amplified in NEB Stable Competent *E. coli* (New England Biolabs #c3040i) or other *E. coli* strains provided by vendors, grown in LB broth (MSKCC Media Core) overnight and isolated with ZymoPURE II kit (Zymo Research #D4203). Mouse cancer cell lines generated in this study were engineered to constitutively express V5-tagged Firefly luciferase (pLenti-PGK-V5-Luc-Puro^w543–1^, Addgene #19360), kind gift from Dr. Eric Campeau and Dr. Paul Kaufman. Some LeptoM derivatives (LLC LeptoM, E0771 LeptoM, B16 LeptoM) used in the flow cytometry experiments were additionally engineered to constitutively express AmCyan fluorescent protein (pLV-EF1a-AmCyan1-IRES-Puro, Takara #0039VCT). Lentiviral constructs for CRISPR-Cas9 editing in the pLV-hCas9:T2A:Bsd backbone were synthetized by VectorBuilder. sgRNA sequences expressed under the control of U6 promoter were as follows: sg*LacZ* - ‘TGCGAATACGCCCACGCGAT’, sg *Ifngr2*#1 ‘TGGACCTCCGAAAAACATCT’, sg*Ifngr2*#2 ‘AGGGAACCTCACTTCCAAGT’, sg *Ifngr2*#3 ‘TCTGTGATGTCCGTACAGTT’. Lentiviral particles were prepared with LentiX 293T cell line using ecotropic, VSV-G pseudotyped lentiviral system and concentrator (Takara #631276 and #631232), as recommended. Mouse cancer cell lines were spin-transduced (1000 g, 32°C, 1 hour) with concentrated lentiviral particles in complete culture medium containing 5 μg/mL hexadimethrine bromide (Santa Cruz, #sc-134220) and selected for 5-7 days in complete medium containing 2-5 μg/mL puromycin (Gibco, #A1113802) or 5- 10 μg/mL blasticidin (Invivogen, ant-bl-1). CRISPR-Cas9 edited lines and control clones were single-cell sorted into 96-well plate. Gene function was assessed functionally (LLC, E0771, and B16 LeptoM; see fig. S8), and DNA editing was confirmed with Sanger sequencing (LLC and E0771 LeptoM; not shown) after expansion.

### Cancer cell injections

Cancer cells were injected into mice between 6 and 16 weeks of age. Mice were deeply anesthetized in an insulated chamber perfused with 2-3% isoflurane (Covetrus; #11695067772) in medical air or with intraperitoneally delivered mixture of ketamine (100 mg/kg) and xylazine (10 mg/kg) in ultra-pure, sterile, and pyrogen-free water for injection. Female mice were used for breast cancer models and both males and females in approximately 1:1 ratio for melanoma and lung cancer models, if not stated otherwise. Mice deceased within 72 hours of injection were excluded from further analysis. Mouse hair was removed from the injection site, and the area was sterilized three times with ethanol. For intracisternal injection, 10 μL of cancer cell suspension in PBS was introduced into the cisterna magna using Hamilton syringe (Hamilton #HT80501) fitted with a 30G needle, as described previously with minor modifications ^13^. Briefly, mouse was positioned prone over a 15 mL conical tube to place cervical spine in flexion. The occiput was palpated, the needle was advanced 4 mm deep, and the syringe content was slowly released into the cisterna magna. The syringe was then held in this position for another ten seconds and then carefully ejected to prevent the reflux of injected liquid. This procedure was tolerated well by the animals (success and survival rate > 95%). Mice displaying neurologic symptoms upon awakening were immediately euthanized. The number of cancer cells introduced intracisternally was: 2,000 cells for LLC LeptoM, 4,000 cells for E0771 LeptoM, and 500 cells for B16 LeptoM, Yumm5.2 LeptoM, EMT6 LeptoM, and 4T1 LeptoM cells. For intracardiac injections, 10,000 cells (for 4T1 or EMT6 sublines) or 50,000 cells (all other sublines) was injected in 50 μL saline using 28G insulin syringe into the left cardiac ventricle. For extracranial injections, cells were injected in 50 μL percutaneously into the fourth mammary fat pad (E0771 LeptoM; 500,000 cells), subcutaneously (LLC LeptoM; 200,000 cells), or intradermally (B16 LeptoM; 100,000 cells) using 28G insulin syringe.

### Quantification of tumor burden

The spread and growth of cancer cell lines engineered to express V5-tagged Firefly luciferase (lucV5) was monitored using non-invasive bioluminescent imaging (BLI). Mice were anesthetized in an insulated chamber perfused with 2-3% isoflurane in medical air and injected retro-orbitally with 50 μL of sterile D-luciferin (15 mg/mL, Goldbio #LUCK-5G) solution in PBS. BLI was captured using IVIS Spectrum-CT (Perkin Elmer). Data were recorded and processed with Living Image (v4.7.2) software. Recorded images were quantified as cranial radiance. For the rare occasion when mice on C57Bl/6 background (without tyrosinase mutation) developed melanin spots preventing luciferase imaging, these animals were not included in the imaging analysis. Tumors in the mammary fat pad, intradermal and subcutaneous tumors were measured with calibrated digital calipers (VWR #62379-531). Tumor volumes are expressed as the product of the two largest diameters, as in ^14^.

### Quantification of leptomeningeal tumor burden with image analysis

B16 melanoma sublines growing in 3D structures produce high amounts melanin that quenches light in a wide spectrum of wavelengths, interfering with accurate bioluminescent and fluorescent imaging. For these tumors, bioluminescence was therefore used solely to confirm the presence or identify the anatomic location of cancer. To overcome this limitation and to accurately quantify the tumor burden in B16 LeptoM model, brains from intracisternally injected mice were dissected, preserving the plaques of cancer, and fixed in formalin overnight. Brains were then carefully washed with tap water and placed into 6-well dishes in 70% ethanol. Brightfield images of fixed brains (basilar plane) were taken using Lumar Stereoscope (Zeiss) against dark background. Data were processed with Fiji/ImageJ (v2.0.0, NIH) as follows: images were converted to 8-bit, each brain was manually encircled, and its area was recorded. The threshold for plaque measurement was first estimated in a small cohort to capture only the plaque areas, and then applied to all subsequent measurements. Percentage of the area of cancer plaques covering the basilar surface of the brain was calculated as the area of plaques divided by the area of brain and multiplied by 100. Since the 8-bit images were monochromatic, this method showed to be robust and reproducible throughout different measurements. Five control brains from mice without cancer, collected for different purposes, were measured and the area of darker structures above the pre-set threshold was less than 1% using this method.

### Derivation of leptomeningeal and parenchymal metastatic cell lines

#### BrM cell lines (brain parenchyma-tropic)

50,000 parental cells were injected intracardially. Hematogenous dissemination was confirmed with BLI approximately 1 hour after injection. Upon confirmation of brain colonization with BLI and development of late-stage cancer symptoms, mice were re-injected with luciferin and euthanized. Brains were dissected and imaged *ex vivo* to confirm colonization of parenchyma. Brains with overt lesions were minced, mechanically dissociated using GentleMACS (Miltenyi Biotec) and digested in a mixture of collagenase (100 U/mL, Worthington #LS005273) and DNAse I (10 U/mL, Worthington #LS006333) in HG DME for 1 hour at 37°C, mechanically dissociated every 20 minutes. Suspension was then washed, filtered through a 70-micron mesh, and seeded into corresponding complete culture media, in which P/S was replaced with Primocin. The medium was changed every day for three days, then every other day. Growing cancer cell colonies were expanded for three passages and named BrM1. These cells were then again injected intracardially and the whole procedure was repeated, leading to the establishment of BrM2 cell lines, competent to colonize brain parenchyma after hematogenous dissemination.

#### LeptoM cell lines (leptomeninges-tropic)

2,000 lucV5-expressing parental cancer cells in 10 μL saline were injected intracisternally. Presence in the CSF was confirmed with BLI approximately one hour after injection. Mice were monitored weekly using BLI and daily checked for the presence of pathophysiological symptoms. When these mice developed neurologic symptoms (moribund behavior, head tilt, seizures, overall weakness) and cancer presence in the CSF was indicated by BLI, luciferin was injected retro-orbitally, and mice were euthanized. Brain was dissected as described in and basilar side of brains as well as basilar meninges of mouse were assessed with BLI *post mortem*. The cranial cavity and brain surface were then washed with approximately 3 mL of saline. This volume was collected, pelleted, resuspended in complete media containing Primocin and maintained as described above for BrM cells. This procedure was repeated once for melanomas or three times for epithelial cancers, leading to the establishment of Inter cell lines. These Inter cells were then injected intracardially and mice were monitored with BLI and treated as described above. Successfully expanded cancer cells that were isolated from these intracardially injected mice were capable to grow colonize leptomeninges and growth in the CSF, hence named LeptoM cells. Three to five biological independent sublines were successfully established *per* cell lines. For transcriptomic analyses, these replicates were processed separately retaining the ID of founder mice. For further *in vitro* and *in vivo* manipulations, these replicates were pooled (in one-to-one *etc*. ratios) and maintained under sub-confluent conditions *in vitro* for limited number of passages (less than 12).

### RNA collection and extraction, and transcriptomic analysis

Cancer cell lines were collected 24 hours after initial seeding of approximately 1x10^6^ cells per 100 mm plate by direct lysis with RLT buffer (Qiagen, component of RNeasy kits). RNA from cell lines was isolated with RNeasy Plus Mini Kit (Qiagen #74136), and sequenced and analyzed as described in ^15^. Resulting HTSeq ^16^ matrices from bulk transcriptome were processed in R Studio with DESeq2 ^17^. Data from LLC cell lines was retrieved from NCBI GEO GSE83132. Newly generated raw and pre-processed data are available through NCBI GEO under accession number GSE221358.

### Collection of mouse CSF and leptomeningeal immune cells

Mice were deeply anesthetized using ketamine/xylazine and transcardially perfused with sterile, ice-cold PBS. Mice were positioned as described in ‘Cancer cell injections’ section, and CSF was collected through the cisternal puncture into the PBS-flushed syringe fitted with a 30G needle. Approximately 15 μL of CSF was collected from each single mouse using this procedure. Blood-contaminated samples were discarded. CSF was flash-frozen on dry ice and stored at -80°C until analysis; or diluted in 200 μL of 4% methanol-free paraformaldehyde (Electron Microscopy Sciences #15714-S) and spun onto microscopic slides to produce cytospins. These were then left to air-dry and stained with hematoxylin QS (Vector Biolabs #H-3404-100). Leptomeningeal immune cells were collected as described previously ^1^ and processed further for downstream applications, as described in corresponding sections.

### Intracisternal delivery of recombinant proteins and AAV particles

Vehicle (PBS), or a 10 ng or 25 ng dose of recombinant mouse IFN-**γ** (Biolegend #714006) in total volume of 10 μL was initially delivered with cancer cell injection, followed by weekly administration, as described above. Heat inactivated IFN-**γ** was prepared by incubating vehicle or vehicle-diluted IFN-**γ** at 95°C for 15 min and allowed to cool on ice before administration. Mouse *Ifng* [NM_008337.4] or *Egfp* sequences were inserted into AAV expression vector (pscAAV backbone under the control of CMV promoter) and used for packaging into AAV5 particles that were ultra-purified for *in vivo* applications (VectorBuilder). Genomic content (GC) was estimated with PCR. 5 μL of vehicle-diluted AAV5 suspension (1x10^13^ GC/mL) was slowly infused into mouse leptomeninges intracisternally and mice were allowed to rest for at least 2 weeks before further manipulation.

### Mouse single-cell proteogenomics

#### Sample processing

*Cx3cr1*^GFP/GFP^*Ccr2*^RFP/RFP^ mice were crossed with wild-type C56Bl/6 mice and the resulting female and male *Cx3cr1*^+/GFP^*Ccr2*^+/RFP^ progeny was intracisternally infused with AAV and LLC LeptoM cancer cells, as described above and in fig. S15. Leptomeningeal cells from six animals *per* group were isolated and resuspended in Cell Staining Buffer (Biolegend #420201). In total, we profile leptomeningeal immune cells from 24 mice and 4 different conditions. To limit the non-specific antibody binding, cells from each mouse were incubated with TruStain FcX (Biolegend #101320), subsequently barcoded with TotalSeq-A anti-mouse hashtags 1 to 6 (Biolegend), listed in table S4, and washed. Cells from these six mice were then pooled, resulting in four independent samples, and stained with a custom TotalSeq-A panel (Biolegend) consisting of 198 antibodies targeting cell surface epitopes and non-targeting isotype controls, listed in table S5, to facilitate identification and origin of selected immune cell types (such as in fig. S17, H and I). Dead cells and debris were removed with LeviCell (LevitasBio), washed cells were counted, encapsulated with Chromium Single Cell 3’ GEM Library and Gel Bead Kit V3.1 (10x Genomics), and sequenced on NovaSeq 6000. Quality control plots are shown in fig. S15. Raw and pre-processed data are available through NCBI GEO under accession number GSE221593. Code for pre-processing and downstream analysis is available from corresponding author and will be deposited to GitHub after peer review.

#### Data preprocessing and initial processing

Raw FASTQC files were pre-processed with SEQC ^2, 3^ with modified mouse reference genome mm10 that included GFP, RFP and AmCyan sequences, and pre-processed as human samples, with the exception that no batch correction was applied. Each sample was processed separately. Cell filtering and doublet removal with DoubletDetection (p_thresh=1e-16, voter_thresh=0.5, n_iters=25, use_phenograph=False, standard_scaling=True) ^4^ was performed as described above for human samples, we initially detected 54,781 cells and 20,804 genes and retained 46,852 high-quality cells and 18,277 genes after filtering out low quality cells and non-immune cell populations. We detected ∼1,387±866 genes per cell, ∼4,374±5,483 gene counts per cells, out of which 3.15±2.62% were mitochondrial genes (values represent mean ± one standard deviation). Shannon entropy for this uncorrected mouse dataset was computed as described above for human data. AnnData files for each sample were then merged after filtering and doublet removal by an outer join. Erythrocyte genes (HBA-A1, HBB-BT, HBA-A2, HBB-BS, ALAS2, HBB-BT, HP, and BPGM) and CD41 protein signal (platelet marker) were filtered out, in an addition to mitochondrial (prefix MT-) and ribosomal genes (prefix RPS- or RPL-). HTO and CITE-seq data were demultiplexed with cite-seq-count ^18^, using default parameters applied on the whitelist of cells that passed the filtering step based on RNA quality, as described above. RNA and protein data (HTO and CITE) were integrated with totalVI, facilitating identification of immune cell subtypes using both gene and surface protein expression (default settings with top 4,000 HVG) ^19^. HTOs were assigned based on maximum number of observed counts (as shown in fig. S15E). UMAP kNN graph and Leiden clustering ^6^ in this dataset was computed using sc.pp.neighbors ^10^ and totalVI processed latent variables. Leiden clusters were merged according to major cell types, which were assigned based on marker gene and surface protein expression, as showed in fig. S3. (HVG - highly variable genes).

#### Subsetting of cells for downstream analyses, plotting and visualization

Plotting was performed using Scanpy (UMAP, tSNE, heatmaps) ^10^ and scVelo (UMAP, tSNE; this package was not used to infer RNA velocity) ^11^. Embedding density was computed with sc.tl.embedding_density (Fig. 2C). Cell cycle prediction was adapted from tl.score_genes_cell_cycle (Fig. 4F and 5C) ^11^. Subsetting was performed by selecting cell clusters from major populations, shown in Fig. 2B. We included cells from all four conditions, shown in fig. S15: cells isolated from naïve, vehicle-injected or LLC LeptoM-injected animals that were overexpressing *Egfp* (control gene) or *Ifng* specifically in the leptomeninges. For analysis of dendritic cells (DC), ‘cDC’ and ‘pDC’ clusters were subsetted, these cells were expressing CD11c (pan-DC marker) on cell surface. For analysis of natural killer cells (NK), ‘NK’ and ‘Proliferative T/NK’ clusters were subsetted to ensure proper representation of all NK cells. These cells were reclustered with Leiden (resolution = 0.7), and clusters expressing CD3 and TCRβ cell surface markers were excluded, retaining only bona fide NK cells, characterized as Nk1.1^+^ CD3^-^ TCRý^-^. For analysis of both cell types, we run Palantir (default settings - n_components = 5, knn = 30) that allowed us to (i) compute diffusion components, used for tSNE re-embeddings and (ii) access MAGIC-imputed (Markov affinity-based graph imputation of cells) cell counts (Fig. 4C, 5A, and 5D) ^8, 9^. These imputed cell counts were used only for visualization with 2D plots. tSNE plots were re-fitted using multiscale coordinates that are based on diffusion components obtained with Palantir (n_components=5, knn=30). Subsetted DCs were refitted onto tSNE using Palantir multiscale coordinates and annotated with initial Leiden loadings to identify four typical dendritic cell populations. We considered both gene expression data (shown as a heatmap in fig. S16A) and cell surface signals: cDC1 cells are Xcr1^+^, cDC2 cells are CD11b^+^, pDC cells are B220^+^, while CCR7+ cells express *CCR7* gene (Fig. 4B and S17D). Subsetted NK cells were refitted onto tSNE using Palantir multiscale coordinates and re-clustered with Leiden (resolution = 0.3), that resulted in identification of four putative cell states. Naïve NK cells expressed high cell surface levels of CD62L (encoded by *SELL* gene), while activated and proliferative cells had low CD62L levels. Proliferative cells also expressed genes associated with cell cycling, such as *MKI67*, *TOP2A*, and *HMGB2*. Senescent cells expressed CD55 and KLGR1 on their cell surface (Fig. 5B). Cancer cells, characterized by the expression of keratin genes and *CD63*, were subsetted as ‘cancer’ cluster and visualized with UMAP without re-embedding. Cancer cell gene signatures were computed with GSEApy (fig. S17, B and C; cut-offs are provided in corresponding figure legends) (https://github.com/zqfang/GSEApy).

#### Trajectory analysis

To predict the maturation trajectories of conventional dendritic cells in normal, non-perturbed steady-state mouse leptomeninges and leptomeninges with metastasis, we subsetted CD11c-positive cDC cells from naïve and cancer-bearing mice overexpressing Egfp only (‘cDC’ cluster and ‘egfp’ condition). We first used CellRank to identify putative trajectories without the need for initial or terminal state selection ^20^. We filtered out genes present in less than 10 cells, normalized counts *per* cell and with *log*(X+1) and extracted HVGs with Scanpy’s functions sc.pp.filter_genes, sc.pp.normalize_total, sc.pp.log1p, and sc.pp.highly_variable_genes. We retained 2,635 cells and 2,090 cDC-expressed HVG. We recomputed PCA with sc.pp.pca (n_comps = 50, zero_centered = True) and refitted the tSNE plot with top 9 diffusion components in multiscale space (n_components=9, knn=15), this tSNE map was used for further visualization. We used cytoTRACE kernel ^21^ that allowed us to assess plausible and biologically traceable cell transitions, following their trajectory from more primitive to mature cells. We imputed gene counts from normalized and filtered count matrix with scv.pp.moments with default parameters (n_pcs = 30 , n_neighbors = 30) and initialized CellRank’s cytoTRACEkernel with default parameters. Transition matrix was computed (threshold_scheme = hard). Given that this approach provides qualitative insights into the transition matrix by iteratively choosing the next cell based on the current cell’s transition probabilities, we further compared two additional settings: (i) we did not specifiy from which cells or condition to select starting point (start_ixs = None), or (ii) we selected all cells from naïve *Egfp*-overexpressing mouse as the starting points. Both approaches identified CCR7+ DCs as mature endpoints, and to remain agnostic to the initiation, we continued the analysis without initial cells or states being defined (n_sim = 100). We used GPCCA estimator (Generalized Perron Cluster Cluster Analysis) ^22^ to coarse-grain a discrete Markov chain into a set of macrostates, and compute coarse-grained transition probabilities among the macrostates. We identified three macrostates and assigned each cell their dominant microstate membership. These results suggested that the cDC2 population is prone to maturate towards CCR7+ DCs, with insignificant contribution of cDC1 population (fig. S17, F to H). CellRank prediction was corroborated by analysis with Palantir (n_components = 9, knn = 15, num_waypoints = 500) ^9^ that identified cDC2 population as the one with the highest entropy (maturation potential), and this observation was robust to change in the number of diffusion components, neighbors, or waypoints (fig. S17, I and J). We dissected the cDC2-to-CCR7 DC transition axis and plotted smoothened gene trends along predicted Palantir pseudotime axis (Fig. 4D).

### Bone marrow chimeras

Male C57Bl6-*Tyr*^c-2^ mice were initially anesthetized with 2-3% isoflurane in medical air and restrained in ventilated conical plastic tubes. Animals were placed in a prone position and irradiated using X-RAD320 irradiator (Precision; North Branford, CT, USA) with the following settings: 250kV; 12mA; using 0.25 mm copper filter; distance of radiation source to the animal body: 50 cm; irradiation field: 20 × 20 cm; dose rate: 117.5 cGy/min. Five animals were fitted into the radiation field and received and two cycles of 5.5 Gy total body radiation 6 hours apart. Immediately after completion of the irradiation procedure, animals were returned to their cages and fed with sulfatrim-enriched diet for the duration of this experiment. Within 24 hours, mice were retro-orbitally infused with approximately 1x10^7^ bone marrow cells from multiple pooled wild-type or *Zbtb46-*DTR^+/+^ C57Bl/6 donors. Bone marrow cells were sterilely isolated from *femur* and *tibia*. Inner bone marrow was exposed and placed inside a 0.6 mL PCR tube with small hole punched in the bottom. The PCR tube was placed in 1.5 mL microcentrifuge tube and the samples were spun down to collect and pellet the bone marrow cells. Cells were counted and resuspended in sterile PBS.

### Immune cell depletions

Monocyte-macrophages were depleted with rat anti-mouse CSF1R antibody (Bio X Cell #BE0213), rat IgG2a isotype was used as control (Bio X Cell #BE0089). Antibodies were diluted in in sterile pH 7.0 (BioXCell #IP0070) and delivered intraperitoneally. An initial dose of 400 μg was injected one day before cancer cell implantation, followed by tri-weekly injection of 200 μg. Monocyte-macrophages were also independently depleted with anionic clodronate liposomes, vehicle-containing liposomes were used as control (Clophosome®-A and Control Liposomes, FormuMax Scientific #F70101C-AC-10). Liposomes were delivered retro-orbitally. Initial dose of 200 μL was injected one day before cancer cell implantation, followed by bi-weekly injection of 100 μL. cDC progenitors were depleted in bone marrow chimers that received wild-type or *Zbtb46-*DTR^+/+^ donor cells with diphteria toxin (DTx; Sigma #D0564), diluted in PBS, and delivered intraperitoneally. Initial dose of 400 ng was injected one day before cancer cell implantation, followed by bi-weekly injection of 100 ng. Both wild-type and *Zbtb46-*DTR^+/+^ cohorts were receiving DTx. NK cells were depleted with polyclonal rabbit anti-mouse asialo GM1 (Poly21460; Biolegend #146002), rabbit polyclonal IgG was used as control (Invitrogen #02-610-2). Both antibodies were reconstituted with PBS. Initial dose of 50 μg was instilled one day before cancer cell implantation, followed by bi-weekly injections of 50 μg.

### Flow cytometry

Single-cell suspensions were prepared as described above and in ^1^. After filtering though 70- micron filter and washing with 2 mM EDTA and 1% BSA in PBS, nonspecific binding sites were blocked with TruStain FcX (Biolegend #101320) diluted in PBS, supplemented with 10% rat serum (Sigma #R9759) for 10 min on ice. Antibodies against surface antigens were diluted in reconstituted Brilliant Stain Buffer Plus (BD #566385), supplemented with 5% rat serum. Surface antigens were stained for 15 min on ice. LIVE/DEAD Green/Violet/FarRed Dead Cell Stain kits (Life Technologies #L34969, L34963, L34973, respectively), DAPI (Molecular Probes #D1306) or propidium iodide (Thermo Fisher #P3566) were used as viability stains. Buffer without BSA was used before LIVE/DEAD staining, which was performed for 15 min on ice. Red blood cells were lysed with 1X ACK buffer or 1x eBioscience RBC Lysis Buffer (Invitrogen #00-4300-54) for 5 min at ambient temperature. For cytokine production analysis, leptomeningeal isolates were resuspended in serum-free IMDM and incubated (MSKCC Media Core) with or without addition of brefeldin A (Biolegend # 420601), ionomycin (StemCell Technologies # 73722), and phorbol 12-myristate 13- acetate (PMA; Invivogen # tlrl-pma), for 2 hours at 37°C. Where the intracellular staining was performed, cells were further fixed with IC Fixation Buffer for 20 min (Invitrogen, 00-8222-49) at room temperature, permeabilized and stained with antibodies against intracellular markers in 1x Permeabilization Buffer for 1 h (Invitrogen, 00-8333-56) and analysed. For pSTAT1 transcription factor staining, cells were processed using True-Nuclear Buffer Set (Biolegend #424401) or FOXP3 Fix/Perm Buffer Set (Biolegend #421403) and analysed. MHC class I levels of vehicle- or recombinant mouse IFN-**γ** -exposed (Biolegend #714006) cells *in vitro* was measured 24 hours after treatment. Data was recorded using LSR Fortessa (BD). Gating and analysis was performed essentially as described in ^1, 23^, using unstained samples, isotype-stained samples, and/or FMO controls. Antibodies used for flow cytometry are listed in table S6.

### Soluble protein detection in plasma and CSF

Solute analytes in the human and mouse CSF were analyzed using following multiplexed bead arrays, used as recommended by the manufacturer: LEGENDPlex mouse anti-virus response (Biolegend #740622), LEGENDPlex human CD8/NK panel (Biolegend #740267).

### NK cell *in vitro* survival assay

NK cells were enriched from dissociated spleens of female and male C57Bl/6 mice with MojoSort mouse NK cell isolation kit (Biolegend #480049). Approximately 20,000 cells were seeded into 1:1 mixture of HG DME and human CSF from cancer patients without LM, containing 10 ng/mL recombinant human IL2 (Biolegend #589102), into 96-well plate. Cells were incubated for 24 h with or without the addition of 1 ng/mL or recombinant mouse IL12p70 (Biolegend #577002) and recombinant mouse IL15 (PeproTech #210-15-10ug). Viability and cell counts were assessed with cytometry.

### Histology

Tissue from euthanized mice was fixed in 10% formalin overnight, thoroughly washed in tap water, sliced, and stored in 70% ethanol until embedded into paraffin. Paraffin-embedded blocks were then cut into 5 micron thick sections and placed onto microscopic slides. Hematoxylin & eosin (H&E) stains were performed by MSKCC Molecular Cytology Core. Myelin stain was performed with Luxol Fast Blue stain kit (Abcam #ab150675). Immunofluorescence was performed as described in ^1^, using following primary antibodies: CD11c (hamster, 1:50, Novus #NBP1-06651 and #NB110-97871, used in combination); Cleaved Caspase 3 (rabbit, 1:200, Cell Signaling Technology #9661S); CNPase (mouse, 1:1000, Abcam #ab6319); DCX (sheep, 1:200, R&D #AF10025); GFAP (goat, 1:500, Abcam #ab53554); Iba1 (rabbit, 1:500, Invitrogen #PA5-27436; and goat, 1:500, Novus #NB100- 1028), MBP (mouse, 1:100, R&D #MAB42282); NeuN (mouse, 1:100-1:500, Sigma #MAB377); Olig2 (goat, 1:200, R&D #AF2418). AF488-, Cy3-, and AF647-conjugated, anti-mouse, goat, rabbit, and sheep secondary antibodies were obtained from Jackson ImmunoResearch; AF647-conjugated anti-hamster secondary antibody was obtained from Abcam. For antibodies of murine origin applied on mouse tissue, the endogenous IgG was first blocked with reconstituted VisUBlock Mouse (R&D #VB001-01ML). DAPI (Molecular Probes #D1306) was used as nuclear counterstain. Autofluorescence was quenched with Vector TrueView (Vector Laboratories #sp-8400). Slides were scanned with Mirax slide scanner (Zeiss), and images for further analysis were exported with CaseViewer (3DHISTECH).

### Quantification of immunofluorescence imaging

Quantification of Iba1+ myeloid cells in choroid plexus was performed essentially as described in ^24^. Cleaved Caspase 3-positive cells in cancer plaques and clusters in leptomeninges were counted manually in FOVs of approximately equal size. Cancer plaques in timepoint-matched AVV5-*Ifng* animals are rare; 2-3 FOVs per brain were extracted and the exact animal sample size and number of FOVs is stated in the corresponding images. Analysis of NeuN was done in the motor and somatosensory cortex in two regions. Region #1 covers layers 1-4 and region #2 covers layer 5 and 6. One section per animal was analysed and the chosen sections were spanning levels -0.18 to -0.196 relative to bregma. Olig2 was quantified in the *corpus callosum*, spanning a lateral area from 0-1.7 mm relative to bregma; and together with CNPase in also in subcortical and cortical region above *corpus callosum*. All image analyses were performed in Fiji/ImageJ ^25^.

### Statistical analysis and reproducibility

Plotting and statistical analysis was performed with Prism 8.1.0 (GraphPad Software), using Mann-Whitney U test, unless specified otherwise. In the box plots (box & whisker plots), box extends from 25^th^ to 75^th^ percentile and whiskers show minimum to maximum values. Results from single-cell analyses were plotted in Python. Bulk RNA-seq was processed in R Studio. All mouse experiments are from at least two independent repetitions, except for fig. S6, C and D; fig. S8, G to I; fig. S9J; fig. S13H; and fig. S20C. Whenever possible, mice were randomly allocated into treatment groups. This was not possible in experiments with transgenic animals. Investigators were not blinded to genotype or treatment over the course of experiment. Sample size and exact *P* values are included in figures. Sample sizes were not pre-determined. Critical mouse experiments (Fig. 2 and 3) were reproduced by two independent investigators. Animal exclusion criteria for animal experiments (death within three days of injection or appearance of pigmented spots that interfered with BLI) are described above. No human samples were excluded from analyses.

---

**Fig. S1.**
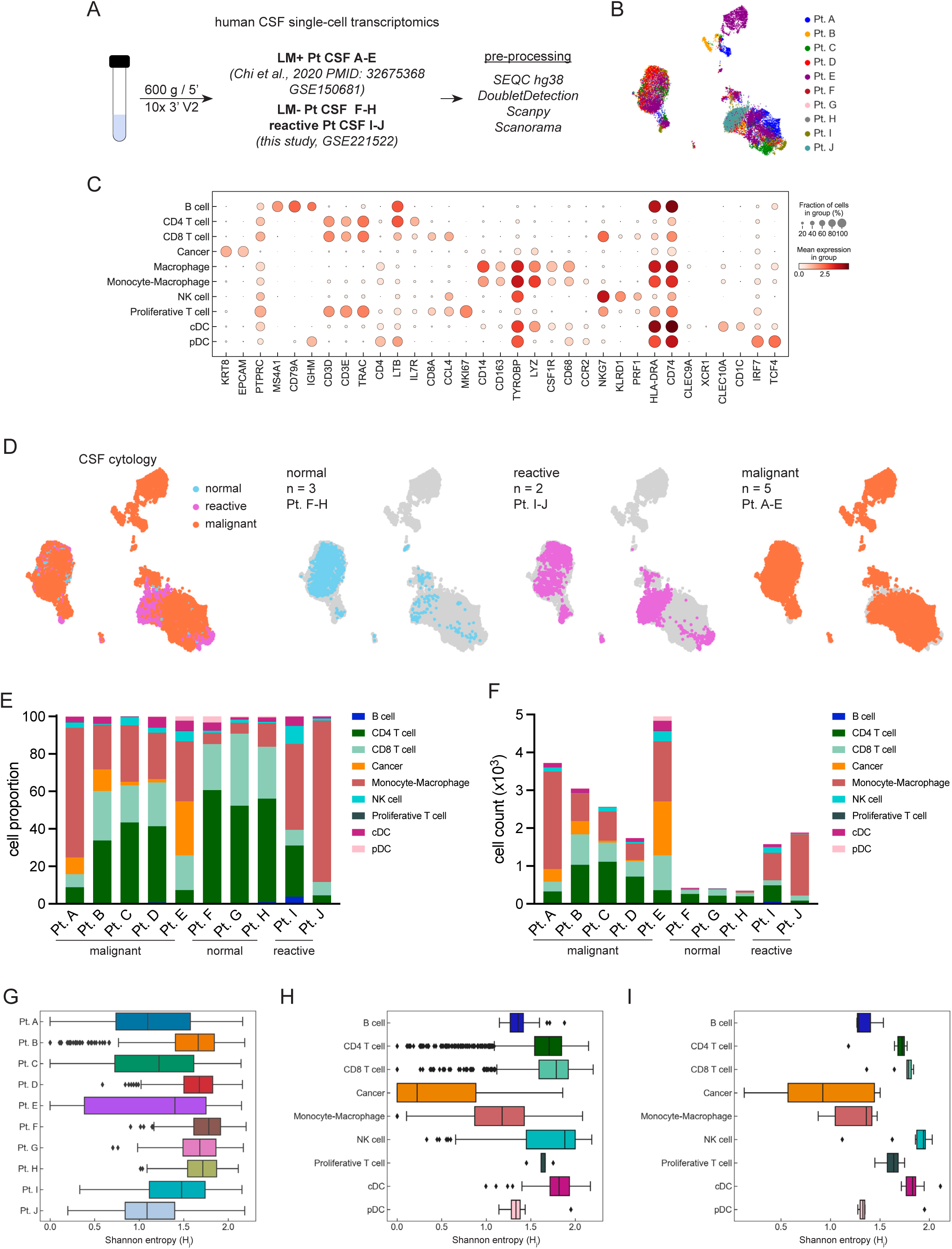
Single-cell transcriptomics of normal, reactive, and malignant human CSF. (**A**) Experimental overview of human CSF single-cell transcriptomics. Single-cell RNA-seq data from five LM+ patients with malignant cytology was retrieved from GSE150660 and integrated with previously unpublished data from three patients with negative cytology and two patients whose CSF contained reactive cells. All samples were processed within the same timeframe. For details, see Methods and panels B to I. (**B**) UMAP of human CSF cells colored by individual patient (n = 20,676 cells from n = 10 patients). (**C**) Expression of cell type-specific marker genes in human CSF single-cell dataset. (**D**) UMAP of human CSF immune cell types and cancer cells grouped based on cytology, and UMAPs of individual cytologies projected separately. Normal: n = 3 patients and n = 1,196 cells; Reactive: n = 2 patients and n = 3,458 cells; Malignant: n = 5 patients and n = 16,022 cells. (**E**) Proportion of major cell types in the individual patients. (**F**) Cell counts of major cell types in the individual patients. (**G**) Inter-patient heterogeneity measured with Shannon entropy in subsampled dataset, where we randomly selected up to 500 cells per patient. For each cell, the Shannon entropy measures the sample diversity of its nearest neighbors in the kNN graph. (**H**) Inter-sample heterogeneity measured with Shannon entropy in subsampled dataset. (**I**) Inter-sample heterogeneity measured with Shannon entropy in subsampled dataset, averaged *per* cell type and *per* patient.

**Fig. S2.**
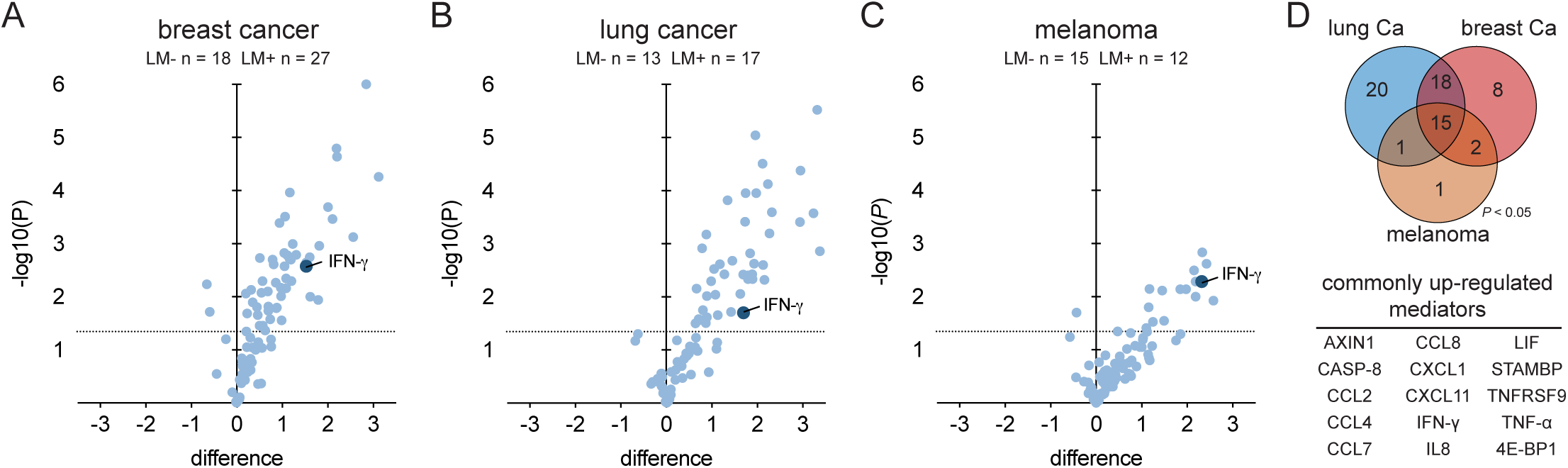
Targeted proteomics with proximity extension assay of inflammatory mediators in human CSF. (**A**) Targeted proteomic analysis of 92 inflammatory mediators in CSF of breast cancer patients without and with LM by proximity extension assay (multiple t tests). (**B**) Targeted proteomic analysis of 92 inflammatory mediators in CSF of lung cancer patients without and with LM by proximity extension assay (multiple t tests). (**C**) Targeted proteomic analysis of 92 inflammatory mediators in CSF of melanoma patients without and with LM by proximity extension assay (multiple t tests). (**D**) Overlap of inflammatory mediators significantly enriched in CSF of LM+ patients, plotted *per* primary cancer type (Venn diagram, top panel). Overview of 15 proteins enriched in CSF from LM+ patients and all three cancer types.

**Fig. S3.**
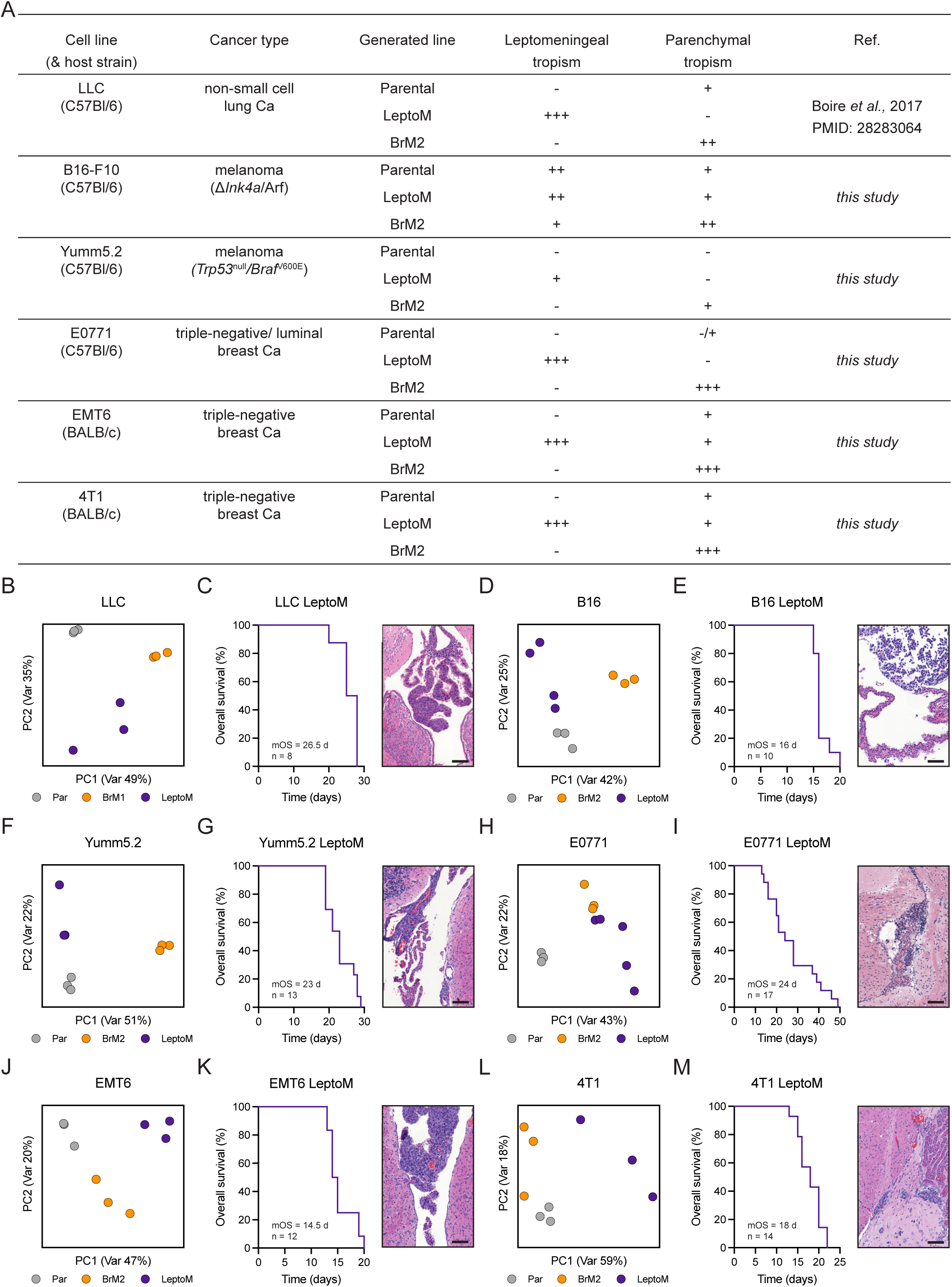
Immunocompetent mouse models of leptomeningeal metastasis. (**A**) Overview of cancer cells lines used and generated in this study. (**B**) Principal component analysis (PCA) of *in vitro* transcriptome of Parental (gray, n = 3), LeptoM (purple, n = 3), and BrM1 (orange, n = 3) LLC cells. Retrieved from NCBI GEO GSE83132. (**C**) Kaplan-Meier plot showing survival of C57Bl/6-*Tyr*^c-2^ animals overexpressing *Egfp* in the leptomeninges after delivery of LLC LeptoM cells into *cisterna magna* (related to fig. S10). Representative brain tissue sections stained with H&E showing colonization of leptomeninges after intracardiac delivery of LLC LeptoM cells (scale bar = 100 μm). mOS - median overall survival. (**D**) Principal component analysis (PCA) of *in vitro* transcriptome of Parental (gray, n = 3) and newly established LeptoM (purple, n = 4), and BrM2 (orange, n = 3) B16 cells. (**E**) Kaplan-Meier plot showing survival of C57Bl/6 animals overexpressing *Egfp* in the leptomeninges after delivery of B16 LeptoM cells into *cisterna magna* (related to fig. S10). Representative brain tissue sections stained with H&E showing colonization of leptomeninges after intracardiac delivery of B16 LeptoM cells (scale bar = 100 μm). mOS - median overall survival. (**F**) Principal component analysis (PCA) of *in vitro* transcriptome of Parental (gray, n = 3) and newly established LeptoM (purple, n = 3), and BrM2 (orange, n = 3) Yumm5.2 cells. (**G**) Kaplan-Meier plot showing survival of C57Bl/6 and C57Bl/6-*Tyr*^c-2^ animals overexpressing *Egfp* in the leptomeninges after delivery of Yumm5.2 LeptoM cells into *cisterna magna* (related to fig. S10). Representative brain tissue sections stained with H&E showing colonization of leptomeninges after intracardiac delivery of Yumm5.2 LeptoM cells (scale bar = 100 μm). mOS - median overall survival. (**H**) Principal component analysis (PCA) of *in vitro* transcriptome of Parental (gray, n = 3) and newly established LeptoM (purple, n = 5), and BrM2 (orange, n = 3) E0771 cells. (**I**) Kaplan-Meier plot showing survival of C57Bl/6-*Tyr*^c-2^ animals overexpressing *Egfp* in the leptomeninges after delivery of E0771 LeptoM cells into *cisterna magna* (related to fig. S10). Representative brain tissue sections stained with H&E showing colonization of leptomeninges after intracardiac delivery of E0771 LeptoM cells (scale bar = 100 μm). mOS - median overall survival. (**J**) Principal component analysis (PCA) of *in vitro* transcriptome of Parental (gray, n = 3) and newly established LeptoM (purple, n = 3), and BrM2 (orange, n = 3) EMT6 cells. (**K**) Kaplan-Meier plot showing survival of BALB/c animals overexpressing *Egfp* in the leptomeninges after delivery of EMT6 LeptoM cells into *cisterna magna* (related to fig. S10). Representative brain tissue sections stained with H&E showing colonization of leptomeninges after intracardiac delivery of EMT6 LeptoM cells (scale bar = 100 μm). mOS - median overall survival. (**L**) Principal component analysis (PCA) of *in vitro* transcriptome of Parental (gray, n = 3) and newly established LeptoM (purple, n = 3), and BrM2 (orange, n = 3) 4T1 cells. (**M**) Kaplan-Meier plot showing survival of BALB/c animals overexpressing *Egfp* in the leptomeninges after delivery of 4T1 LeptoM cells into *cisterna magna* (related to fig. S10). Representative brain tissue sections stained with H&E showing colonization of leptomeninges after intracardiac delivery of 4T1 LeptoM cells (scale bar = 100 μm). mOS - median overall survival.

**Fig. S4.**
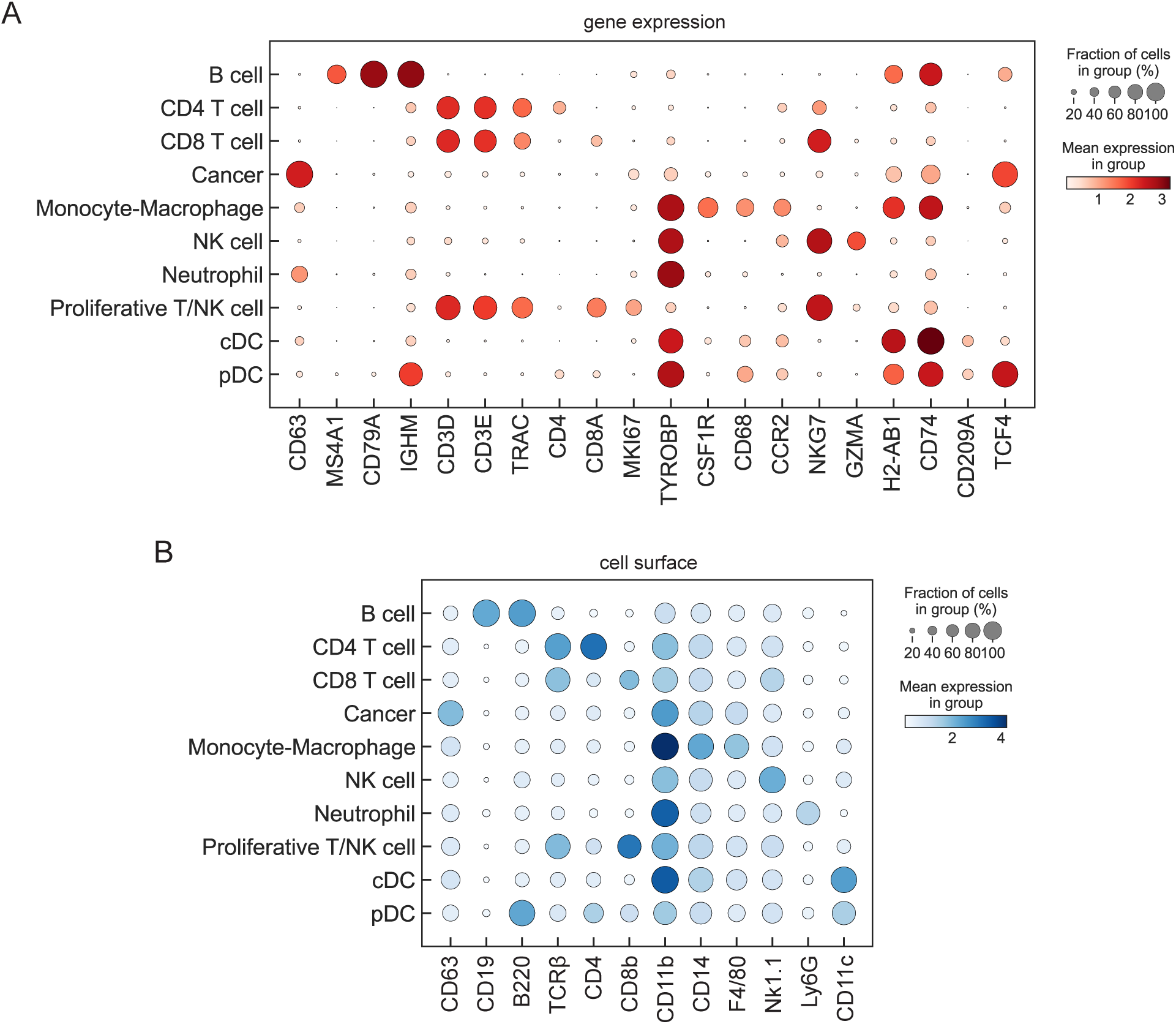
Cell type annotation of mouse leptomeningeal immune cells. (**A**) Expression of cell type-specific marker genes in mouse proteogenomic single-cell dataset, as captured with single-cell RNA-seq. (**B**) Expression of cell type-specific surface markers in mouse proteogenomic single-cell dataset, as determined with CITE-seq.

**Fig. S5.**
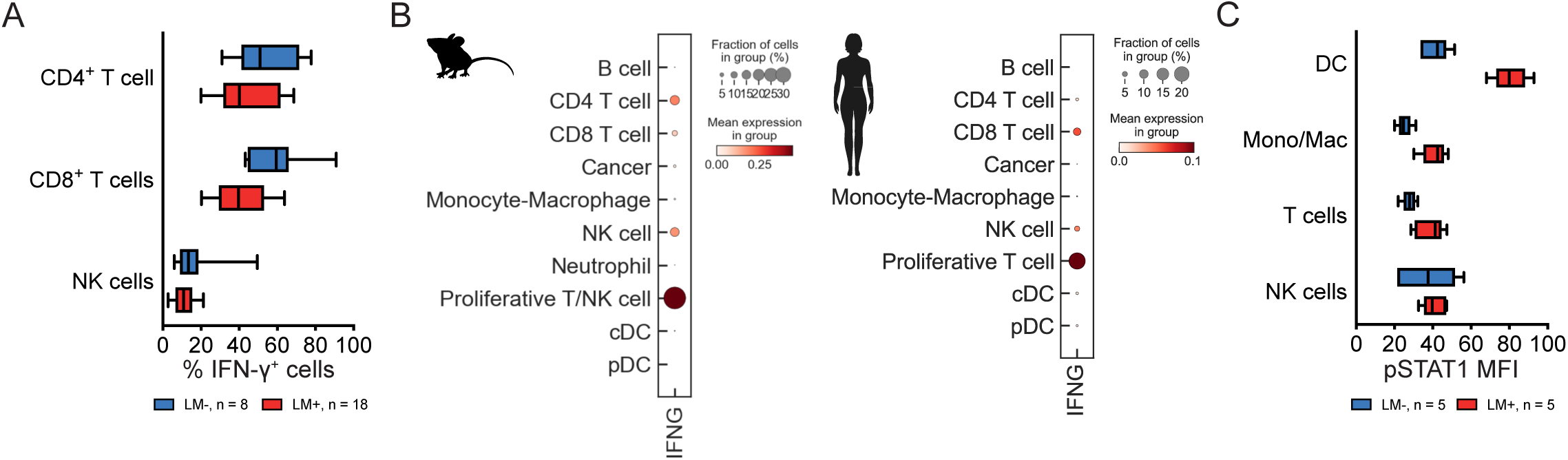
IFN-γ production and response in leptomeninges. (**A**) Proportion of T cells (CD3^+^CD4^+^CD8^-^ vs. CD3^+^CD4^-^CD8^+^) and NK cells (CD3^-^Nk1.1^+^) expressing IFN-**γ** in cells isolated from vehicle- or B16, E0771, and LLC LeptoM-injected mice, determined with flow cytometry. (**B**) Expression of *IFNG* gene in mouse (left) and human (right) single-cell datasets. (**C**) Abundance of phosphorylated STAT1 (pSTAT1) in leptomeningeal dendritic cells (MHC II^+^ CD11c^+^), monocyte-macrophages (CD11b^+^Ly6C^+^ and CD11b^+^F4/80^+^), T cells (CD3^+^), and NK cells (Nk1.1^+^), as a proxy for IFN-**γ** pathway activation in vehicle- and LLC LeptoM-injected mice, determined with flow cytometry.

**Fig. S6.**
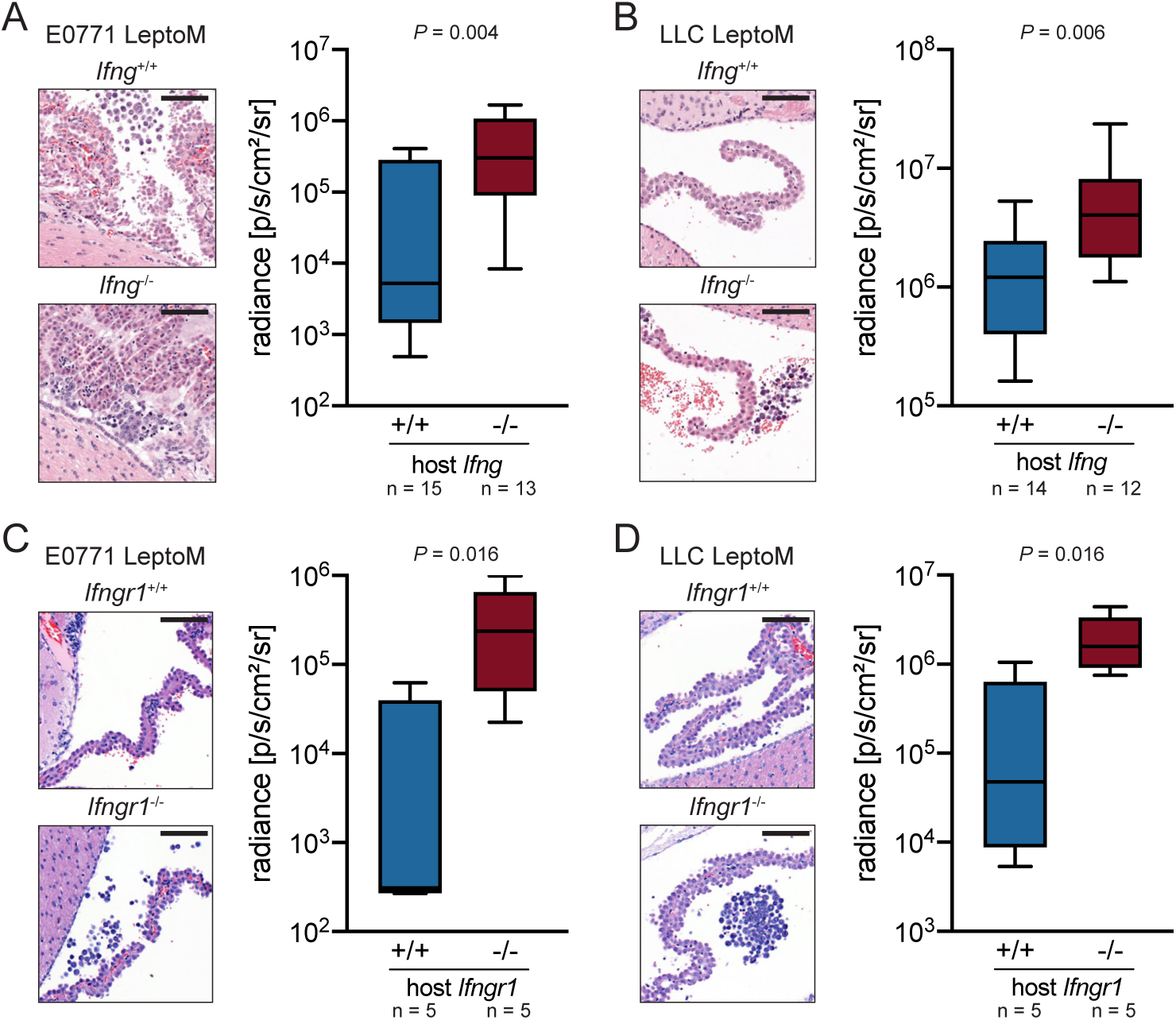
Leptomeningeal tumor growth in *Ifng*- and *Ifngr1*-deficient animals. (**A**) Representative leptomeningeal tissue sections stained with H&E (scale bar = 100 μm). Box plot illustrates *in vivo* radiance of E0771 LeptoM cells delivered intracisternally into C57BL/6 *Ifng*-proficient and -deficient animals, quantified two weeks after injection. (**B**) Representative leptomeningeal tissue sections stained with H&E (scale bar = 100 μm). Box plot illustrates *in vivo* radiance of LLC LeptoM cells delivered intracisternally into C57BL/6 *Ifng*- proficient and -deficient animals, quantified two weeks after injection. (**C**) Representative leptomeningeal tissue sections stained with H&E (scale bar = 100 μm). Box plot illustrates *in vivo* radiance of E0771 LeptoM cells delivered intracisternally into C57BL/6 *Ifngr1*-proficient and -deficient animals, quantified two weeks after injection. (**D**) Representative leptomeningeal tissue sections stained with H&E (scale bar = 100 μm). Box plot illustrates *in vivo* radiance of LLC LeptoM cells delivered intracisternally into C57BL/6 *Ifngr1*-proficient and -deficient animals, quantified two weeks after injection.

**Fig. S7.**
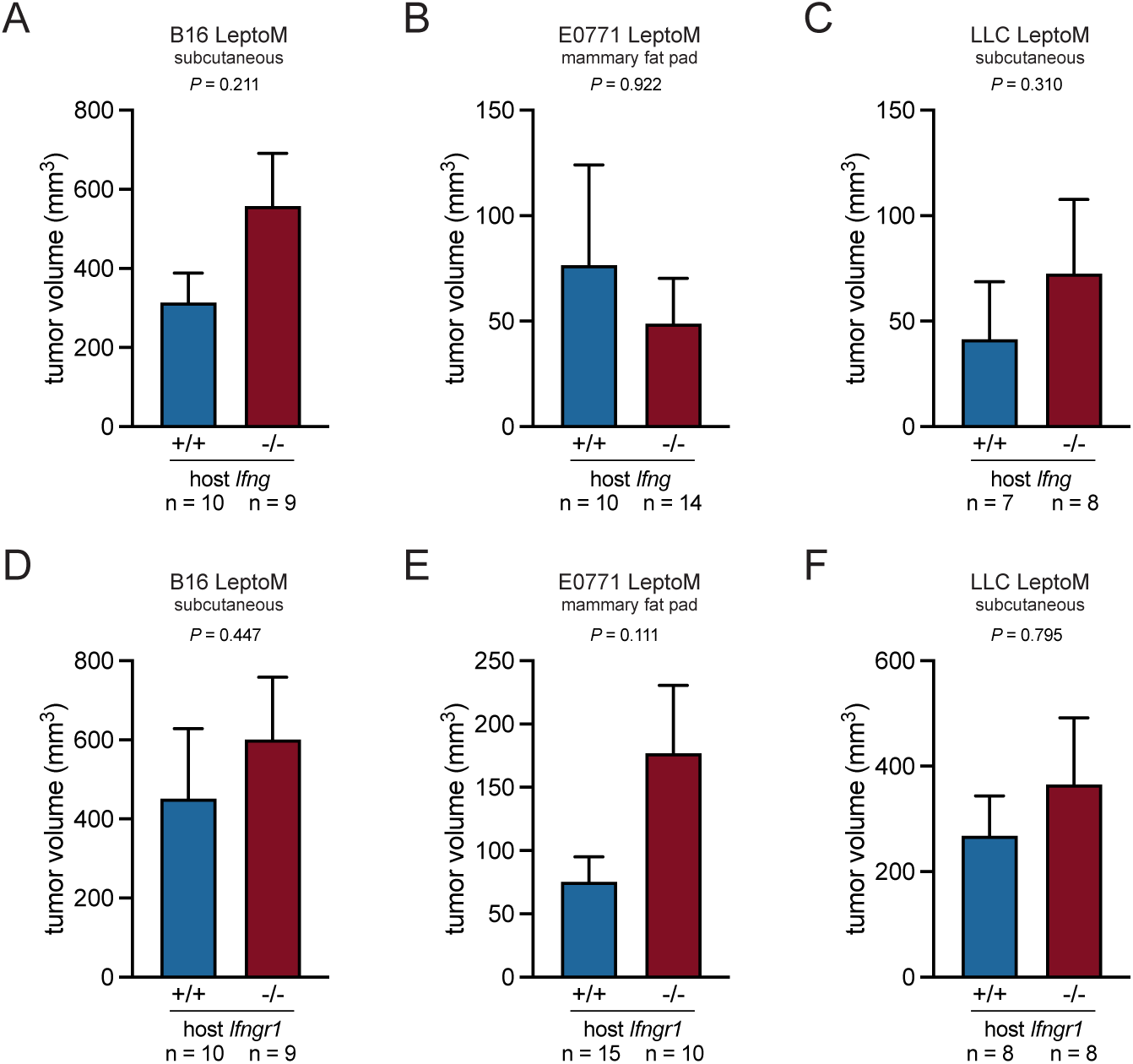
Extracranial tumor growth in *Ifng*- and *Ifngr1*-deficient animals. (**A**) Volumes of intradermal B16 LeptoM flank tumors in C57BL/6 *Ifng*-proficient and -deficient animals, quantified two weeks after injection. (**B**) Volumes of mammary fat pad E0771 LeptoM tumors in C57BL/6 *Ifng*-proficient and - deficient animals, quantified four weeks after injection. (**C**) Volumes of subcutaneous LLC LeptoM flank tumors in C57BL/6 *Ifng*-proficient and -deficient animals, quantified three weeks after injection. (**D**) Volumes of intradermal B16 LeptoM flank tumors in C57BL/6 *Ifngr1*-proficient and -deficient animals, quantified two weeks after injection. (**E**) Volumes of mammary fat pad E0771 LeptoM tumors in C57BL/6 *Ifngr1*-proficient and - deficient animals, quantified four weeks after injection. (**F**) Volumes of subcutaneous LLC LeptoM flank tumors in C57BL/6 *Ifngr1*-proficient and - deficient animals, quantified three weeks after injection.

**Fig. S8.**
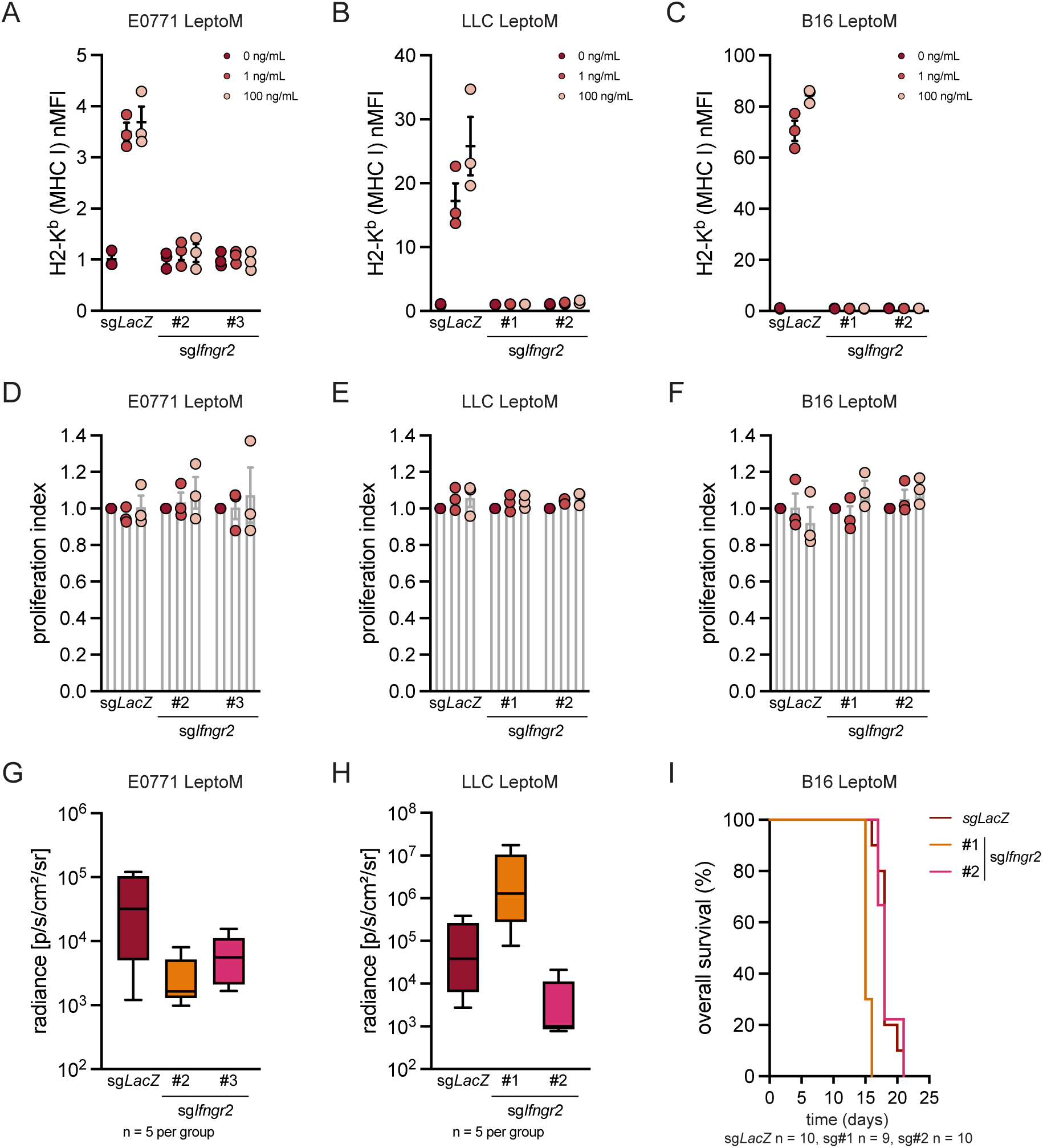
Cancer-intrinsic IFN-γ signaling is dispensable for tumor growth in leptomeninges. (**A**) *In vitro* induction of MHC class I in control (sg*LacZ*) and two *Ifngr2*-deficient E0771 LeptoM clones with recombinant IFN-**γ**. Data pooled from three independent experiments. (**B**) *In vitro* induction of MHC class I in control (sg*LacZ*) and two *Ifngr2*-deficient LLC LeptoM clones with recombinant IFN-**γ**. Data pooled from three independent experiments. (**C**) *In vitro* induction of MHC class I in control (sg*LacZ*) and two *Ifngr2*-deficient B16 LeptoM clones with recombinant IFN-**γ**. Data pooled from three independent experiments. (**D**) *In vitro* proliferation of control (sg*LacZ*) and two *Ifngr2*-deficient E0771 LeptoM clones exposed to recombinant IFN-**γ**. Data pooled from three independent experiments. (**E**) *In vitro* proliferation of control (sg*LacZ*) and two *Ifngr2*-deficient LLC LeptoM clones exposed to recombinant IFN-**γ**. Data pooled from three independent experiments. (**F**) *In vitro* proliferation of control (sg*LacZ*) and two *Ifngr2*-deficient B16 LeptoM clones exposed to recombinant IFN-**γ**. Data pooled from three independent experiments. (**G**) *In vivo* radiance of control (sg*LacZ*) and two *Ifngr2*-deficient E0771 LeptoM clones delivered intracisternally into C57Bl/6-*Tyr*^c-2^ animals, quantified three weeks after injection in one *in vivo* experiment. (**H**) *In vivo* radiance of control (sg*LacZ*) and two *Ifngr2*-deficient LLC LeptoM clones delivered intracisternally into C57Bl/6-*Tyr*^c-2^ animals, quantified two weeks after injection in one *in vivo* experiment. (**I**) Kaplan-Meier plot illustrating overall survival of control (sg*LacZ*) and two *Ifngr2*-deficient B16 LeptoM clones delivered intracisternally into C57Bl/6 mice in one *in vivo* experiment.

**Fig. S9.**
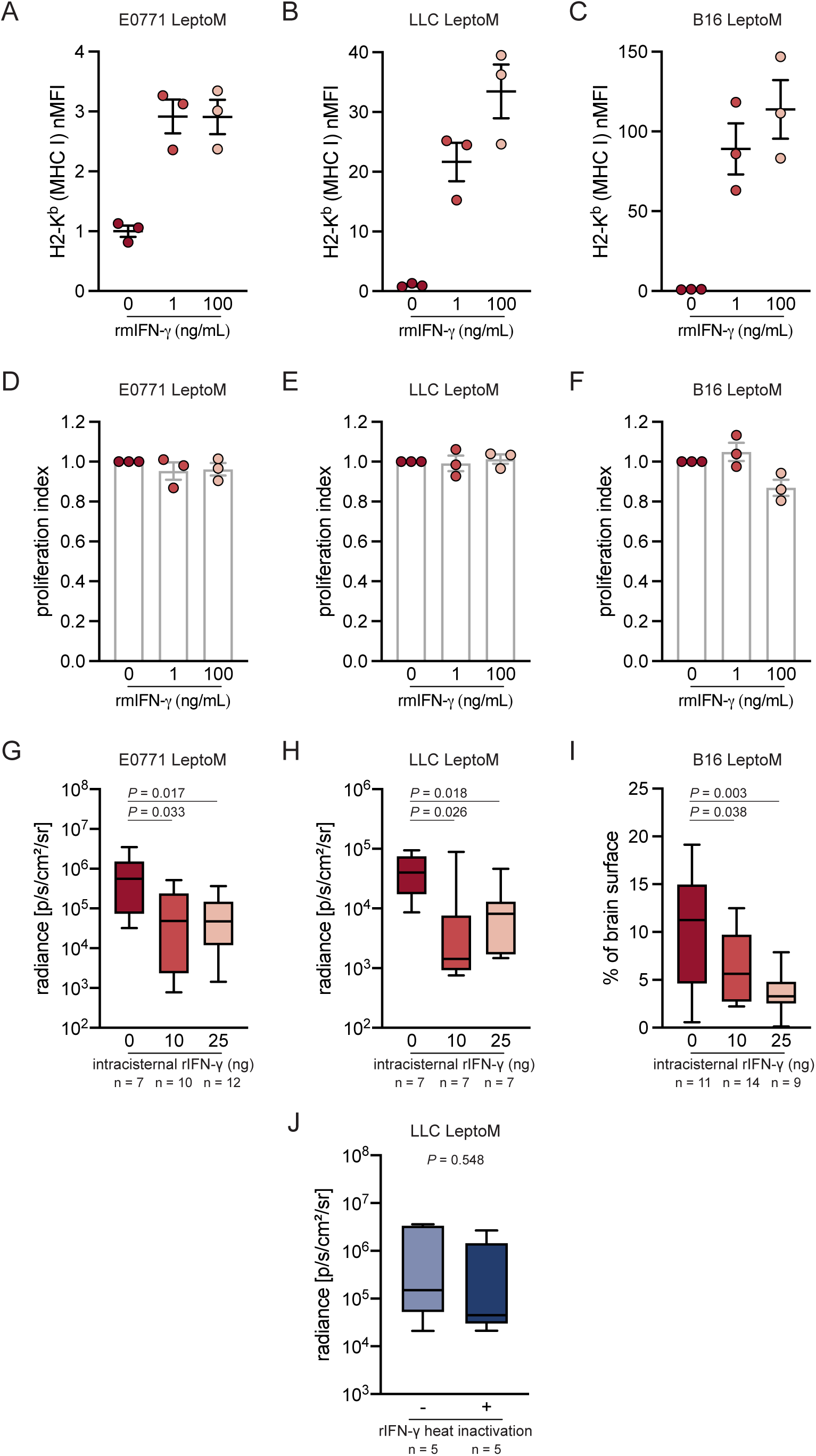
Leptomeningeal IFN-γ -mediated tumor growth suppression is driven by the microenvironment. (**A**) *In vitro* induction of MHC class I in E0771 LeptoM cells with recombinant IFN-**γ**. Data pooled from three independent experiments. (**B**) *In vitro* induction of MHC class I in LLC LeptoM cells with recombinant IFN-**γ**. Data pooled from three independent experiments. (**C**) *In vitro* induction of MHC class I in B16 LeptoM cells with recombinant IFN-**γ**. Data pooled from three independent experiments. (**D**) *In vitro* proliferation of E0771 LeptoM cells exposed to recombinant IFN-**γ**. Data pooled from three independent experiments. (**E**) *In vitro* proliferation of LLC LeptoM cells exposed to recombinant IFN-**γ**. Data pooled from three independent experiments. (**F**) *In vitro* proliferation of B16 LeptoM cells exposed to recombinant IFN-**γ**. Data pooled from three independent experiments. (**G**) *In vivo* tumor growth of E0771 LeptoM cells in C57Bl/6-*Tyr*^c-2^ animals injected weekly with vehicle or two doses of recombinant IFN-**γ**, as a function of radiance. (**H**) *In vivo* tumor growth of LLC LeptoM cells in C57Bl/6-*Tyr*^c-2^ animals injected weekly with vehicle or two doses of recombinant IFN-**γ**, as a function of radiance. (**I**) *In vivo* tumor growth of B16 LeptoM cells in C57Bl/6 animals injected weekly with vehicle or two doses of recombinant IFN-**γ**, as a function of radiance. (**J**) *In vivo* tumor growth of LLC LeptoM cells in C57Bl/6-*Tyr*^c-2^ animals injected weekly with heat-inactivated vehicle (PBS) or heat-inactivated recombinant IFN-**γ**, as a function of radiance in one *in vivo* experiment.

**Fig. S10.**
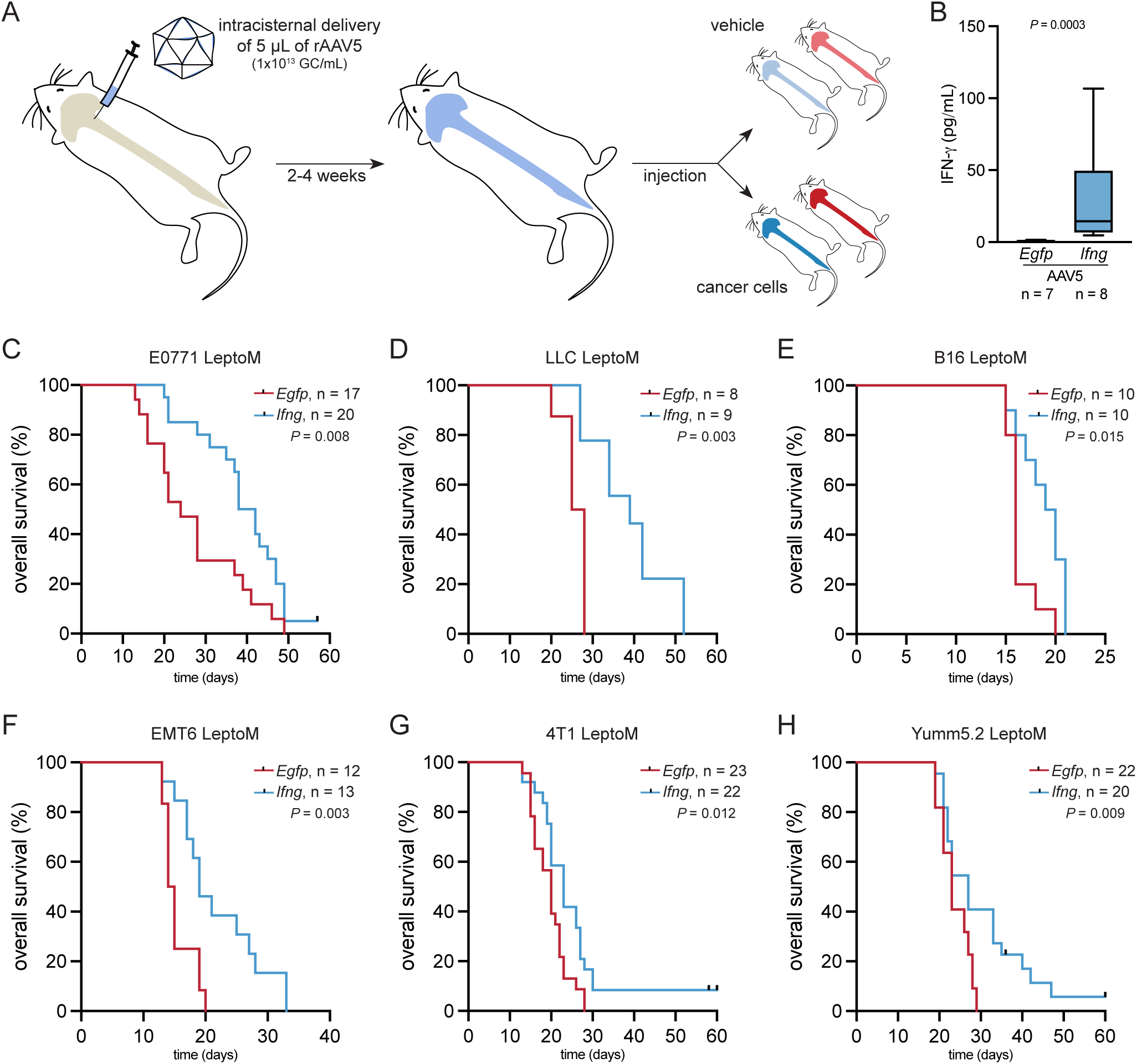
Leptomeninges-specific overexpression of IFN-γ extends survival of LeptoM cells-bearing animals. (**A**) Schematic showing experimental strategy of leptomeningeal *Egfp* or *Ifng* overexpression, used for functional experiments in this study. (**B**) Levels of IFN-**γ** in the CSF collected from naïve C57Bl/6 and BALB/c animals overexpressing *Egfp* or *Ifng* in the leptomeninges, detected by cytometric bead array. (**C**) Kaplan-Meier plot showing survival of E0771 LeptoM-bearing C57Bl/6-*Tyr*^c-2^ animals overexpressing *Egfp* or *Ifng* in the leptomeninges (logrank test). (**D**) Kaplan-Meier plot showing survival of LLC LeptoM-bearing C57Bl/6-*Tyr*^c-2^ animals overexpressing *Egfp* or *Ifng* in the leptomeninges (logrank test). (**E**) Kaplan-Meier plot showing survival of B16 LeptoM-bearing C57Bl/6 animals overexpressing *Egfp* or *Ifng* in the leptomeninges (logrank test). (**F**) Kaplan-Meier plot showing survival of EMT6 LeptoM-bearing BALB/c animals overexpressing *Egfp* or *Ifng* in the leptomeninges (logrank test). (**G**) Kaplan-Meier plot showing survival of 4T1 LeptoM-bearing BALB/c animals overexpressing *Egfp* or *Ifng* in the leptomeninges (logrank test). (**H**) Kaplan-Meier plot showing survival of Yumm5.2 LeptoM-bearing C57Bl/6 and C57Bl/6-*Tyr*^c-2^ animals overexpressing *Egfp* or *Ifng* in the leptomeninges (logrank test).

**Fig. S11.**
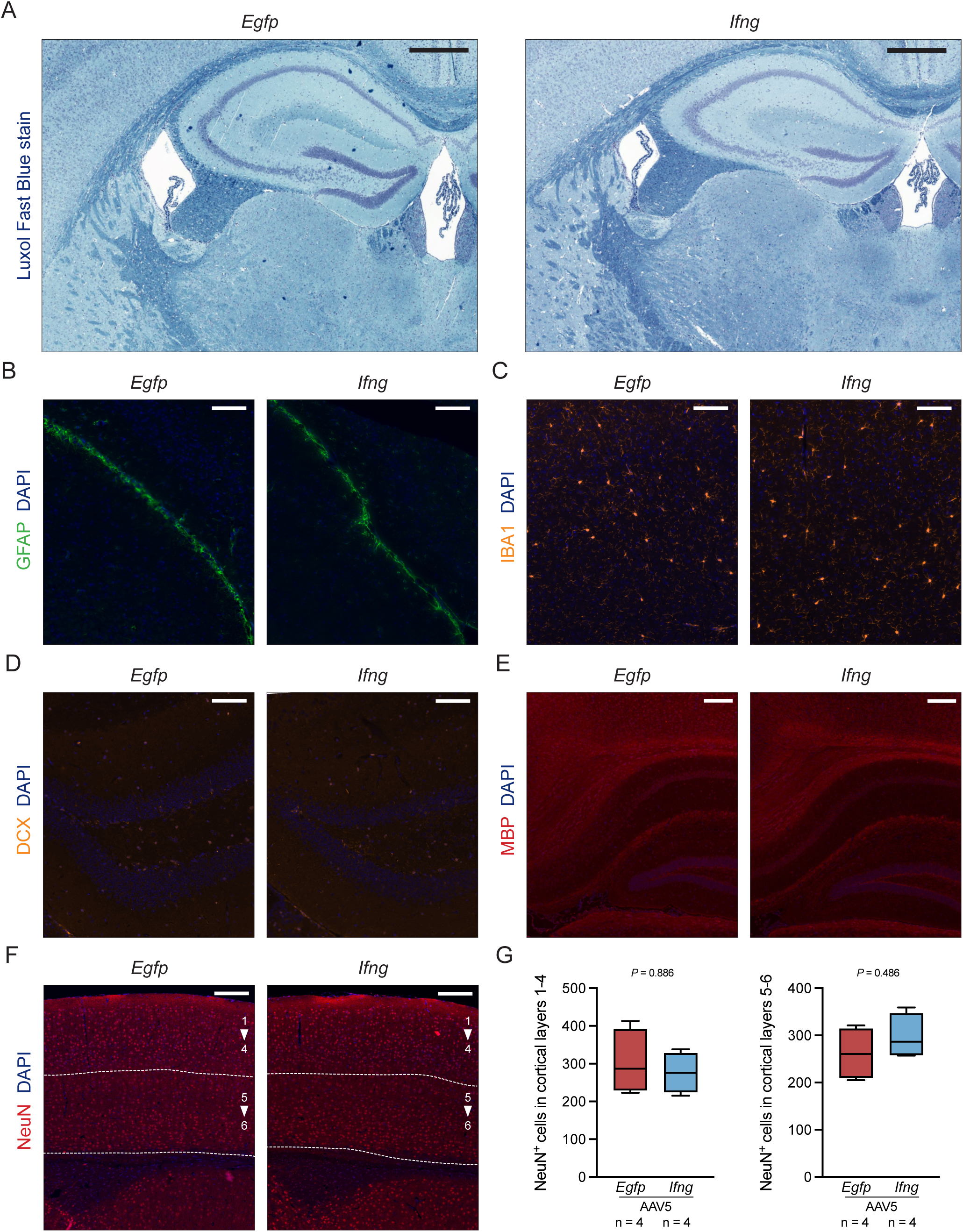
Leptomeningeal IFN-γ does not affect morphology of brain parenchyma. (**A**) Representative images of brain tissue sections from naïve C57Bl/6 animals overexpressing *Egfp* or *Ifng* stained with Luxol Fast Blue (n = 4 *per* group, 3 months after AAV introduction, scale bar = 500 μm). (**B**) Representative images of brain tissue sections from naïve C57Bl/6 animals overexpressing *Egfp* or *Ifng* stained for astrocyte activation marker GFAP (n = 4 *per* group, 3 months after AAV introduction, scale bar = 100 μm). (**C**) Representative images of brain tissue sections from naïve C57Bl/6 animals overexpressing *Egfp* or *Ifng* stained for microglia marker Iba1 (n = 4 *per* group, 3 months after AAV introduction, scale bar = 100 μm). (**D**) Representative images of brain tissue sections from naïve C57Bl/6 animals overexpressing *Egfp* or *Ifng* stained for neural progenitor marker DCX (n = 4 *per* group, 3 months after AAV introduction, scale bar = 100 μm). (**E**) Representative images of brain tissue sections from naïve C57Bl/6 animals overexpressing *Egfp* or *Ifng* stained for myelinization marker MBP (n = 4 *per* group, 3 months after AAV introduction, scale bar = 200 μm). (**F**) Representative images of brain tissue sections from naïve C57Bl/6 animals overexpressing *Egfp* or *Ifng* stained for marker of mature neurons NeuN (n = 4 *per* group, 3 months after AAV introduction, scale bar = 200 μm). (**G**) Quantification of NeuN^+^ mature neurons *per* FOV in cortical layers 1-4 (left) and 5-6 (right). See outline in panel F.

**Fig. S12.**
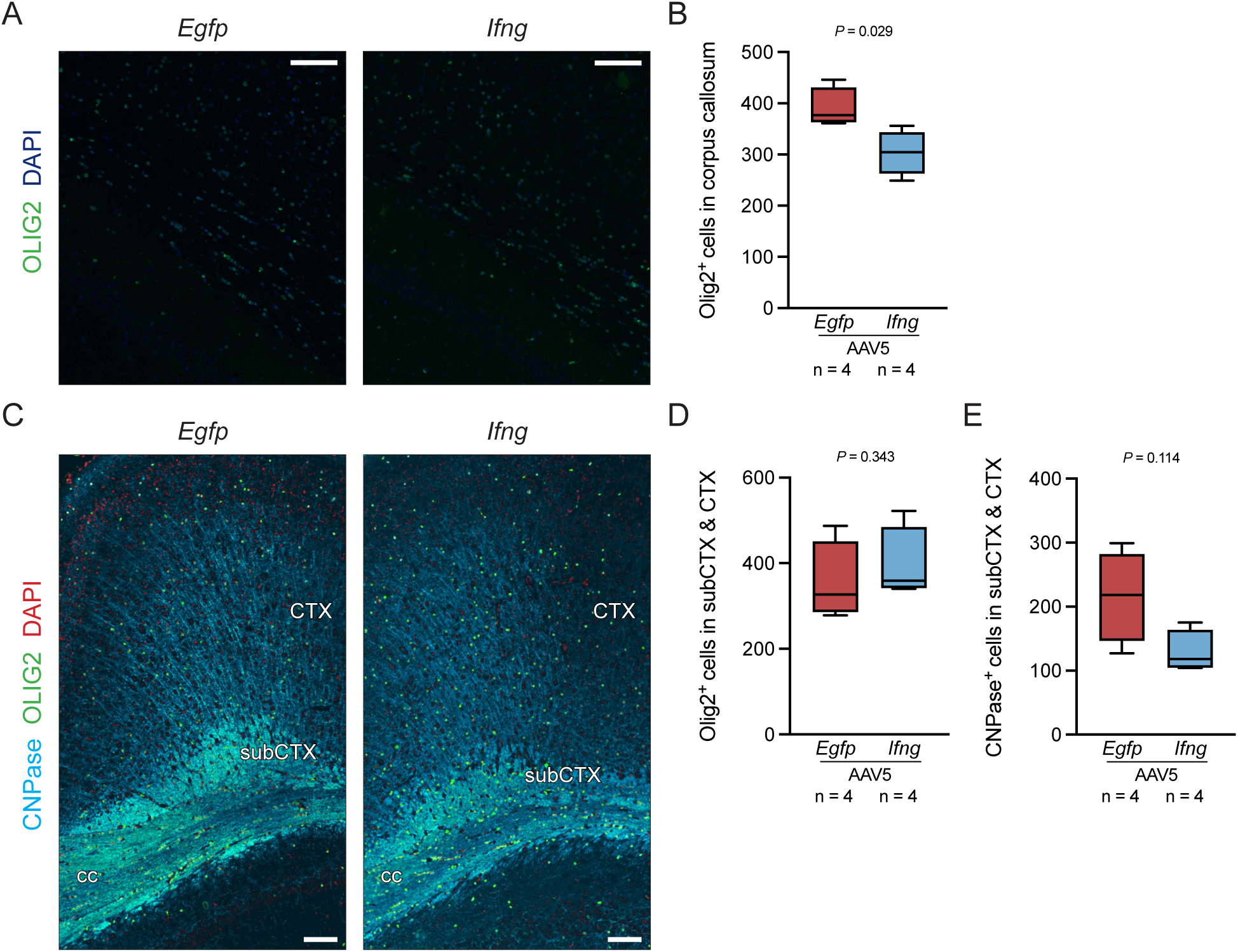
Leptomeningeal IFN-γ reduces oligodendrocyte numbers in *corpus callosum*. (**A**) Representative images of *corpus callosum* sections from naïve C57Bl/6 animals overexpressing *Egfp* or *Ifng* stained for marker of oligodendrocytes Olig2 (n = 4 *per* group, 3 months after AAV introduction, scale bar = 100 μm). (**B**) Quantification of Olig2^+^ oligodendrocytes *per* FOV in *corpus callosum*. (**C**) Representative images of brain tissue sections from naïve C57Bl/6 animals overexpressing *Egfp* or *Ifng* stained for markers of oligodendrocytes Olig2 and CNPase (n = 4 *per* group, 3 months after AAV introduction, scale bar = 100 μm). (**D**) Quantification of Olig2^+^ oligodendrocytes *per* FOV in cortical and subcortical regions. Corresponding regions are marked in panel C. (**E**) Quantification of CNPase^+^ oligodendrocytes *per* FOV in cortical and subcortical regions. Corresponding regions are marked in panel C.

**Fig. S13.**
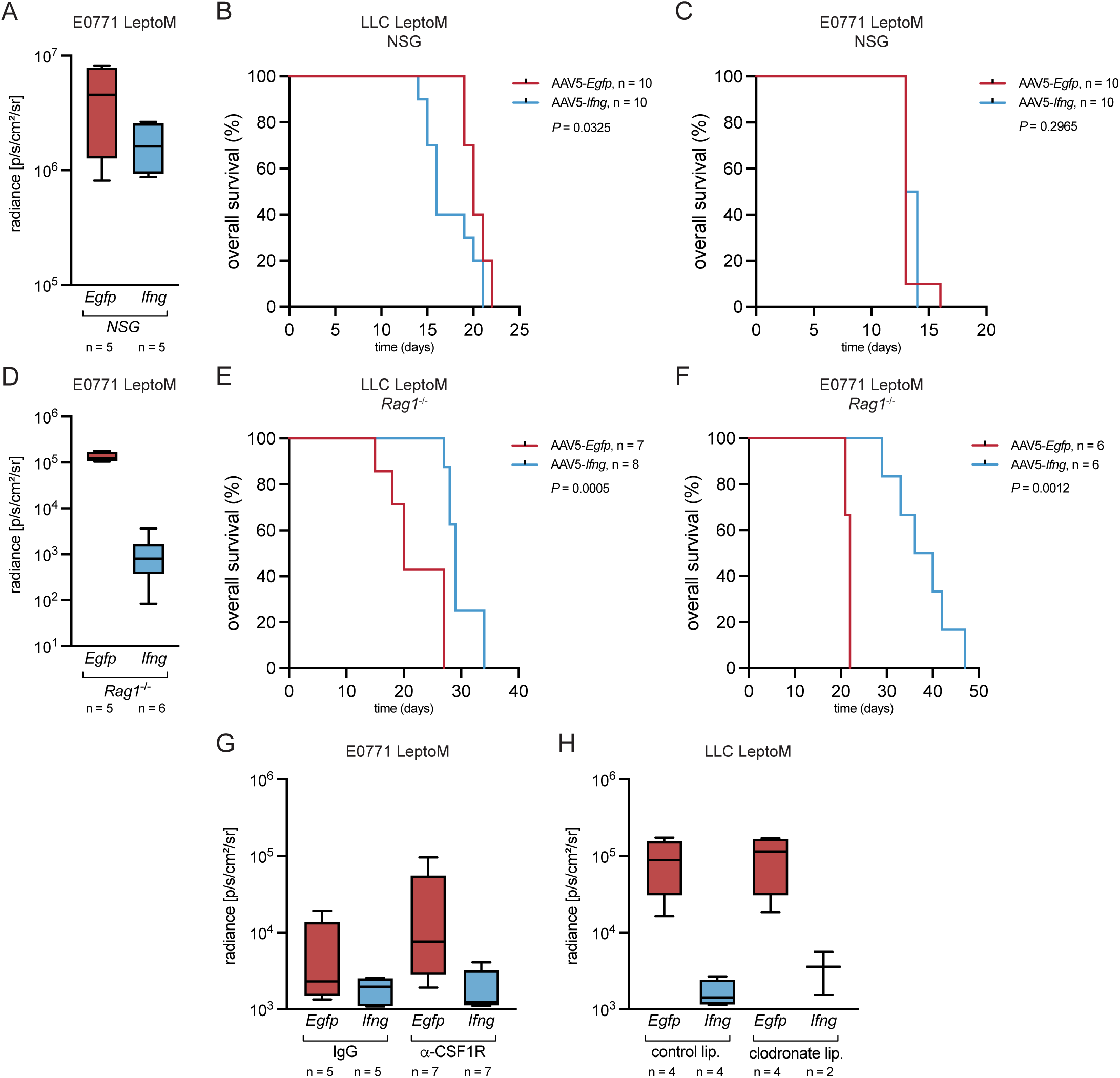
Leptomeningeal IFN-γ does not require adaptive immune system to suppress metastatic outgrowth. (**A**) *In vivo* radiance of E0771 LeptoM cells delivered intracisternally into NSG animals overexpressing *Egfp* or *Ifng* in the leptomeninges, quantified two weeks after injection. (NSG - non-obese, diabetic, severe combined immunodeficient, *Il2rg*^null^). (**B**) Kaplan-Meier plot showing survival of LLC LeptoM-bearing NSG animals overexpressing *Egfp* or *Ifng* in the leptomeninges (logrank test). (**C**) Kaplan-Meier plot showing survival of E0771 LeptoM-bearing NSG animals overexpressing *Egfp* or *Ifng* in the leptomeninges (logrank test). (**D**) I*n vivo* radiance of E0771 LeptoM cells delivered intracisternally into Rag1-deficient animals overexpressing *Egfp* or *Ifng* in the leptomeninges, quantified two weeks after injection. (NSG - non-obese, diabetic, severe combined immunodeficient, *Il2rg*^null^). (**E**) Kaplan-Meier plot showing survival of LLC LeptoM-bearing Rag1-deficient animals overexpressing *Egfp* or *Ifng* in the leptomeninges (logrank test). (**F**) Kaplan-Meier plot showing survival of E0771 LeptoM-bearing Rag1-deficient animals overexpressing *Egfp* or *Ifng* in the leptomeninges (logrank test). (**G**) *In vivo* radiance of E0771 LeptoM cells delivered intracisternally into C57Bl6-*Tyr*^c-2^ animals overexpressing *Egfp* or *Ifng* in the leptomeninges and tri-weekly infused with non-targeting isotype control antibody or CSF1R-targeting antibody, quantified two weeks after injection. (**H**) *In vivo* radiance of LLC LeptoM cells delivered intracisternally into C57Bl6-*Tyr*^c-2^ animals overexpressing *Egfp* or *Ifng* in the leptomeninges and bi-weekly infused with control or clodronate liposomes, quantified two weeks after injection.

**Fig. S14.**
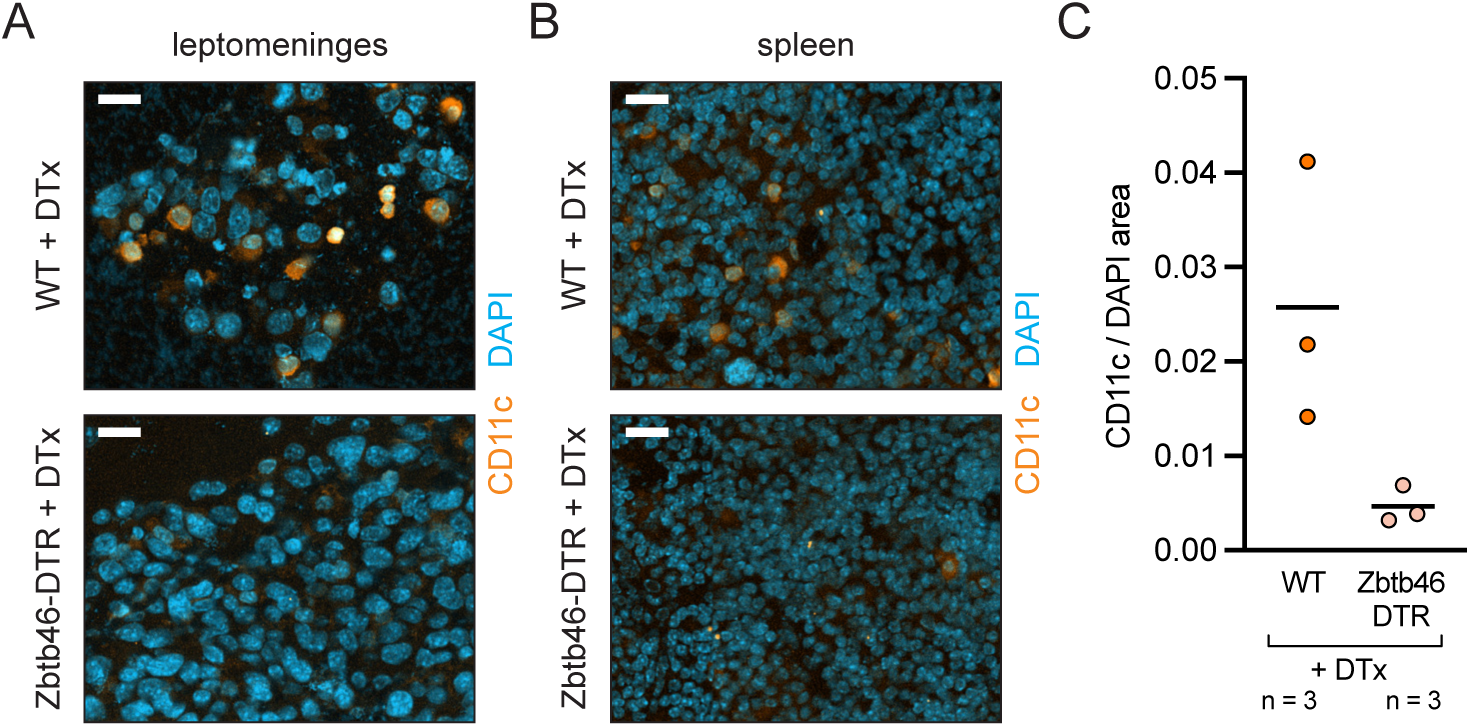
Depletion of leptomeningeal cDCs in bone marrow chimeras. (**A**) Representative images of dendritic cell marker CD11c in leptomeningeal cancer plaques in wild-type (WT) and Zbtb46-DTR bone marrow chimeras, treated with diphteria toxin (DTx). Mice were injected with LLC LeptoM cells (scale bar = 20 μm). (**B**) Representative images of dendritic cell marker CD11c in spleen of wild-type (WT) and Zbtb46-DTR bone marrow chimeras, treated with diphteria toxin (DTx). Mice were injected with LLC LeptoM cells (scale bar = 20 μm). (**C**) Quantification of systemic cDC depletion in images from panel B.

**Fig. S15.**
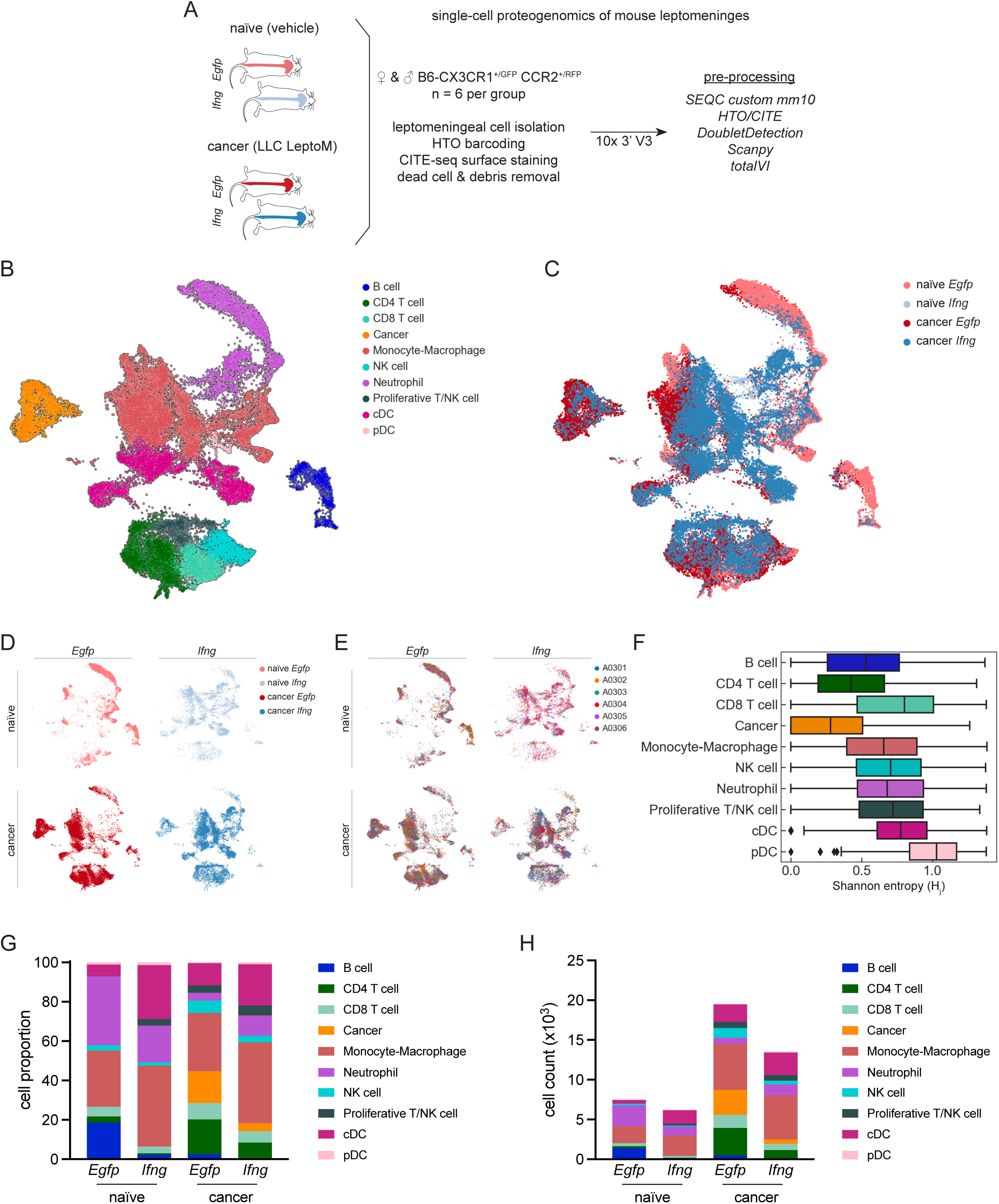
Mouse single-cell proteogenomics of naïve and metastasis-bearing, *Ifng*- overexpressing mice. (**A**) Experimental overview of single-cell proteogenomic analysis of mouse leptomeninges. (**B**) UMAP of mouse leptomeningeal cells colored by major cell type (n = 24). (**C**) UMAP of mouse leptomeningeal cells colored by condition (n = 6 mice *per* group). (**D**) Individual UMAPs of mouse leptomeningeal cells *per* condition. (**E**) Individual UMAPs showing representation of six barcodes *per* condition. (**F**) Inter-sample heterogeneity measured with Shannon entropy. For each cell, the Shannon entropy measures the sample diversity of its nearest neighbors in the kNN graph. (**G**) Proportion of major cell types per condition. (**H**) Counts of major cell types per condition.

**Fig. S16.**
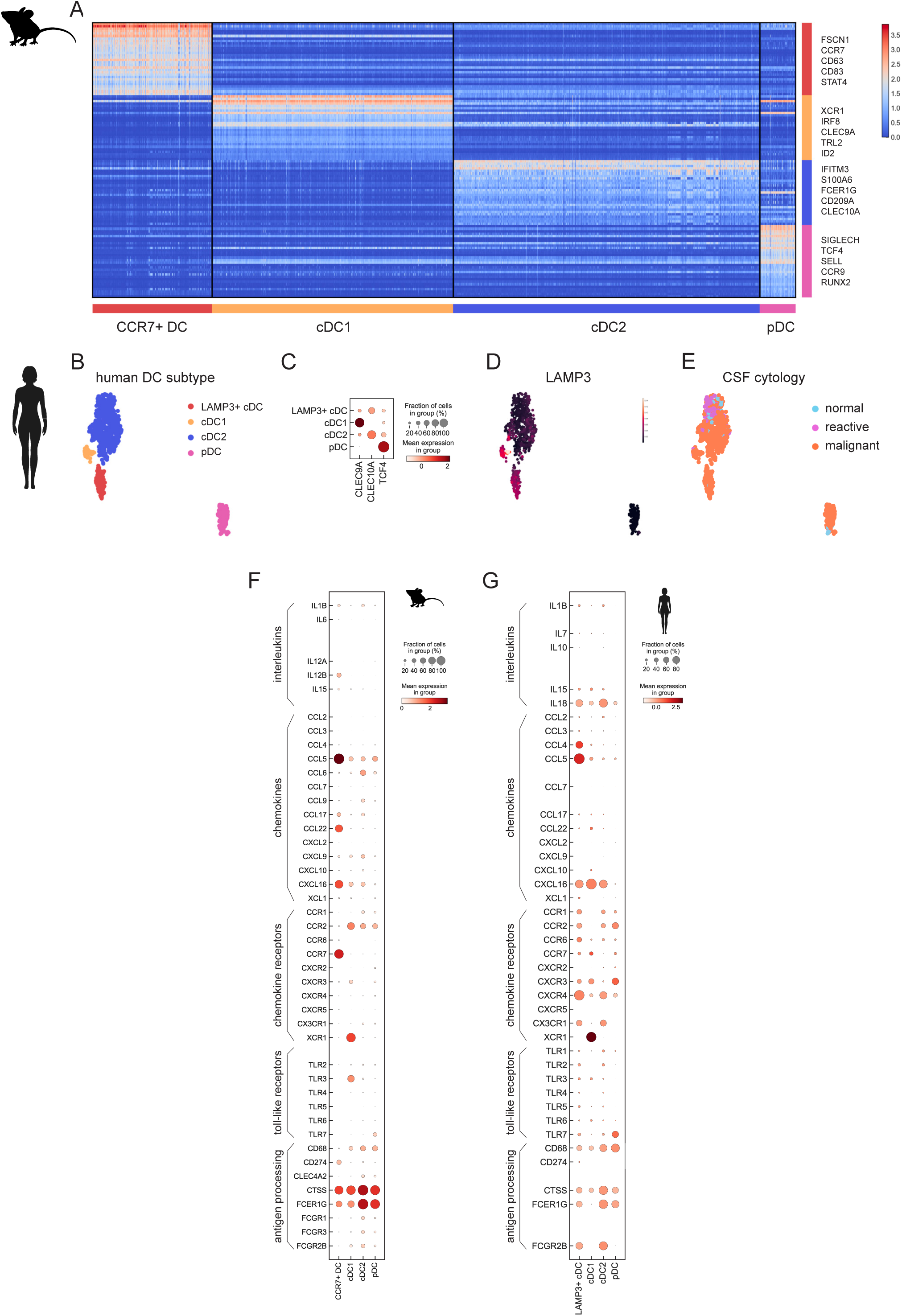
Characterization of leptomeningeal dendritic cells. (**A**) Heatmap showing scaled expression of top 30 genes *per* mouse DC cell type (one cell type vs. the rest; FC > 2). (**B**) UMAP projection of DC subtypes detected in human CSF (n = 883 cells); cDC and pDC clusters from Fig. 1B were subsetted and replotted. cDC1 cells are *CLEC9A*^+^*XCR1*^+^, cDC2 cells are *CLEC10A*^+^*CD1C*^+^, pDC cells are *IRF7*^+^*TCF4*^+^. Human LAMP3+ migratory dendritic cells are *LAMP3*^+^*CCR7*^+^ (orthologous to mouse CCR7+ DC). (**C**) Marker gene expression of human CSF DCs. (**D**) MAGIC-imputed expression of *LAMP3* in human CSF dendritic cells. (**E**) CSF cytology classification of human DC types. (**F**) Dot plot showing gene expression of interleukins, chemokines, chemokine receptors, toll-like receptors, and genes associated with antigen presentation in mouse DC cells, as detected with CITE-seq. Normalized counts were used for computation. Genes not detected with 10x and genes that did not pass filtering steps defined in the Methods were not plotted. (**G**) Dot plot showing gene expression of interleukins, chemokines, chemokine receptors, toll-like receptors, and genes associated with antigen presentation in human DC cells, as detected with scRNA-seq. Normalized counts were used for computation. Genes not detected with 10x and genes that did not pass filtering steps defined in the Methods were not plotted.

**Fig. S17.**
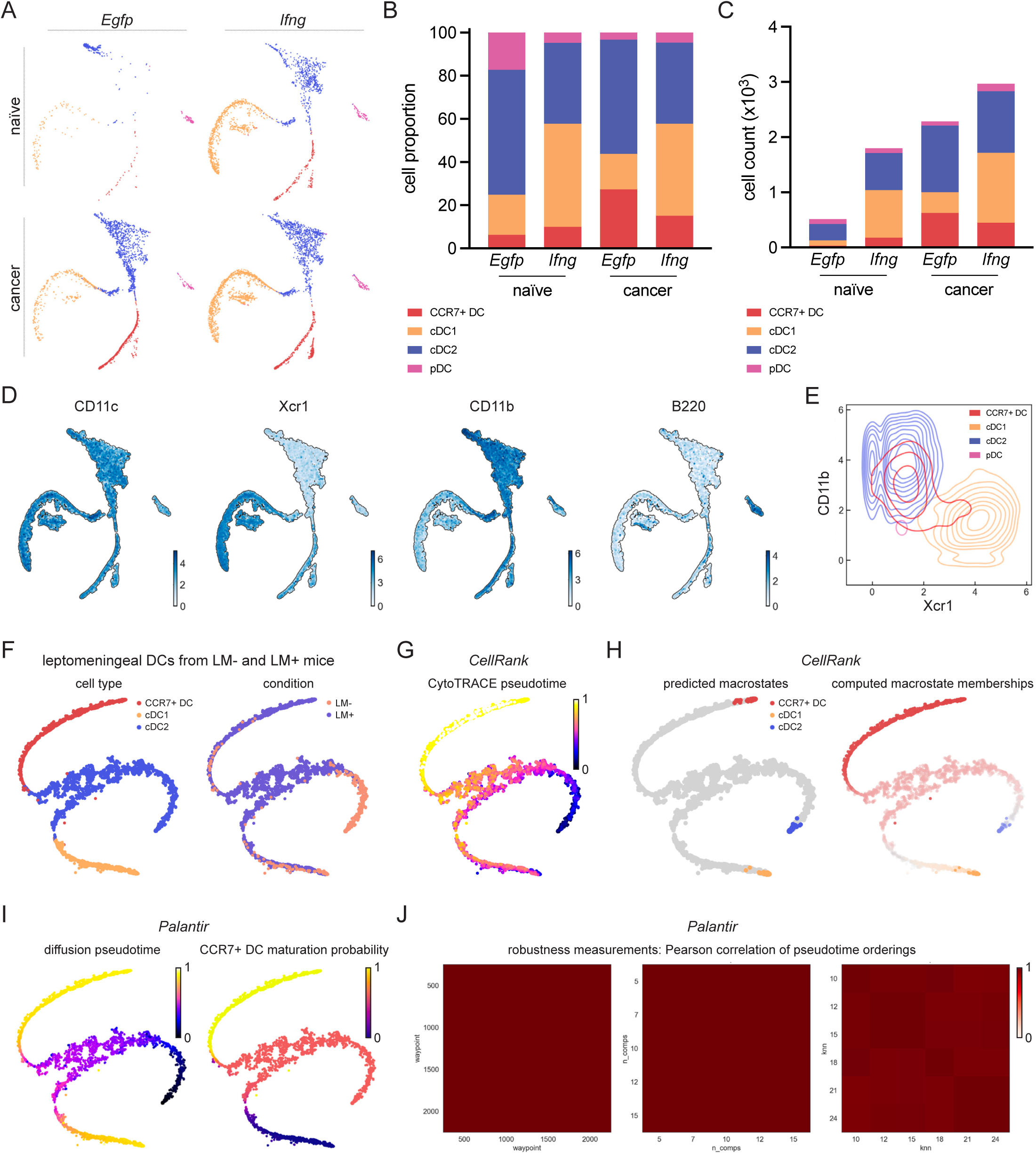
Trajectory analysis of leptomeningeal dendritic cells. (**A**) tSNE maps showing abundance of captured dendritic cell types in naïve and metastasis-bearing, *Egfp*- or *Ifng*-overexpressing mice (total of n = 7,566 cells pooled from 4 conditions and n = 6 animals *per* group). (**B**) Proportion of dendritic cell subtypes per condition. (**C**) Counts of dendritic cell subtypes per condition. (**D**) tSNE projection of dendritic cell surface markers detected with CITE-seq. CD11c - pan-DC marker; Xcr1 - cDC1 marker; CD11b - cDC2 marker; B220 - pDC marker. (**E**) Bivariate plot showing distribution of cell surface Xcr1 and CD11b in leptomeningeal dendritic cell subsets, as detected with CITE-seq. (**F**) tSNE projection of 2,575 mouse leptomeningeal DCs subsetted for trajectory analysis. Cells are from Egfp-overexpressing, naïve and cancer-bearing mice, and the plots are colored based on cell type and condition. See Methods for further details. (**G**) tSNE projection of CytoTRACE pseudotime, as determined with CellRank, suggesting that CCR7+ DCs are the terminal state within the subsetted cell population. (**H**) Terminal DC macrostates and computed macrostate membership for each cell, as predicted with CellRank and projected onto a tSNE. While cDC1 cells are restricted to cDC1 membership, cells from cDC2 cluster are gradually acquiring CCR7+ DC membership. (**I**) Palantir-computed diffusion pseudotime and CCR7+ DC maturation (branch) probability. Gene trends along this pseudotime axis are plotted in Fig. 4D. (**J**) Plots show Pearson correlation of pseudotime orderings in Palantir analysis for different parameters (waypoint samplings, number of principal components, and number of K-nearest neighbours) and all cells. DC trajectory analysis, performed as described in Methods, is not sensitive to fluctuations in these parameters.

**Fig. S18.**
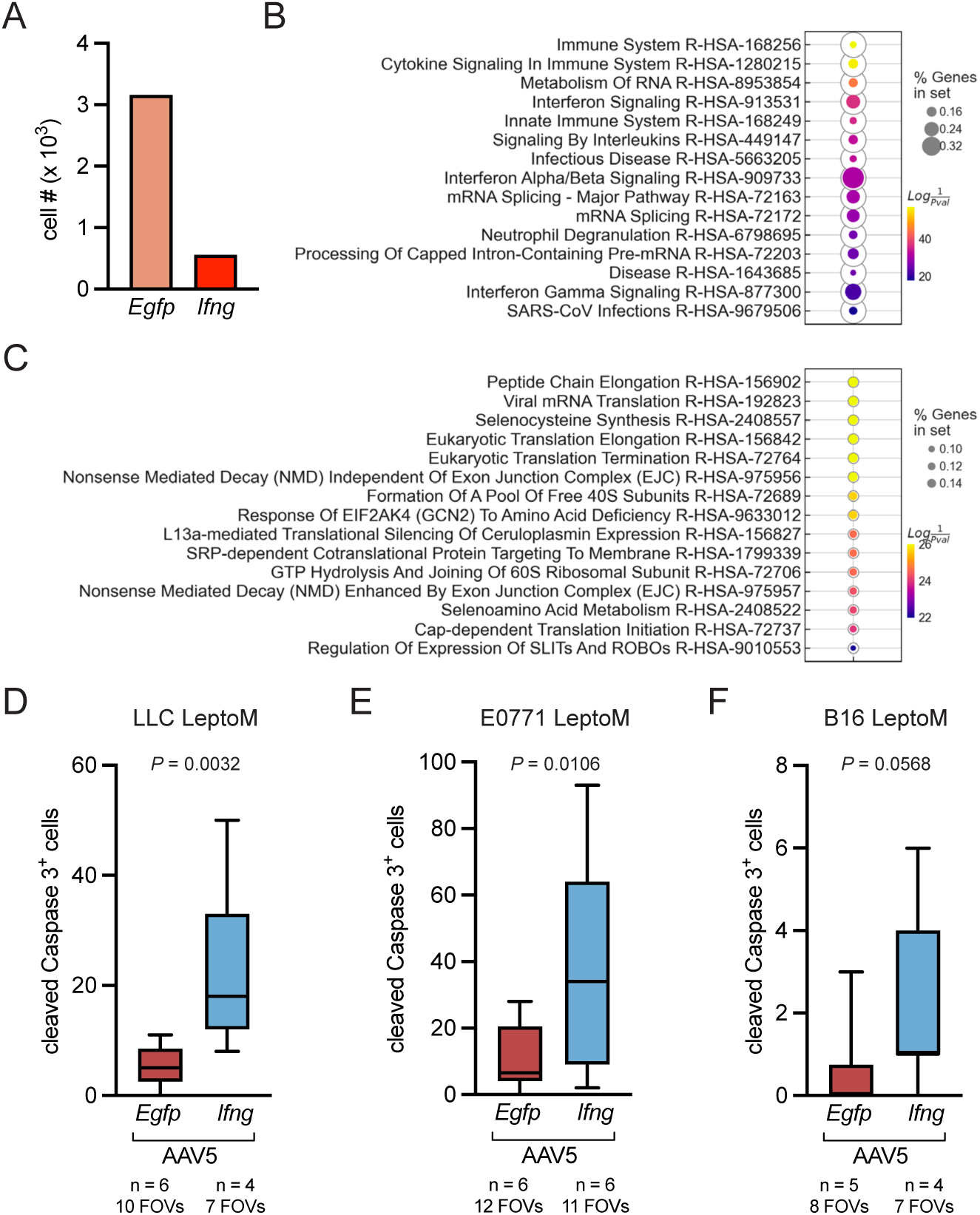
Characterization of leptomeningeal metastatic cells in *Egfp*- and *Ifng*- overexpressing mice. (**A**) Quantification of cancer cells captured in the mouse single-cell atlas (n = 3,718 keratin^+^ CD63^+^ cells isolated from n = 6 mice *per* group); related to Figure 4E, F. (**B**) GSEApy analysis of top 15 Reactome 2022 pathways enriched in cancer cells shown in Fig. 5 isolated from *Ifng*-overexpressing animals and subsetted as described in Fig. 4E (DEG cut-off *P* < 0.01). (**C**) GSEApy analysis of top 15 Reactome 2022 pathways enriched in cancer cells isolated from *Egfp*-overexpressing animals and subsetted as described in Fig. 4E (DEG cut-off *P* < 0.01). (**D**) Quantification of cleaved Caspase 3-positive cells in cancer plaques and clusters, in the leptomeninges of *Egfp*- or *Ifng*-overexpressing animals injected with LLC LeptoM cells. (**E**) Quantification of cleaved Caspase 3-positive cells in cancer plaques and clusters, in the leptomeninges of *Egfp*- or *Ifng*-overexpressing animals injected with E0771 LeptoM cells. (**F**) Quantification of cleaved Caspase 3-positive cells in cancer plaques and clusters, in the leptomeninges of *Egfp*- or *Ifng*-overexpressing animals injected with B16 LeptoM cells.

**Fig. S19.**
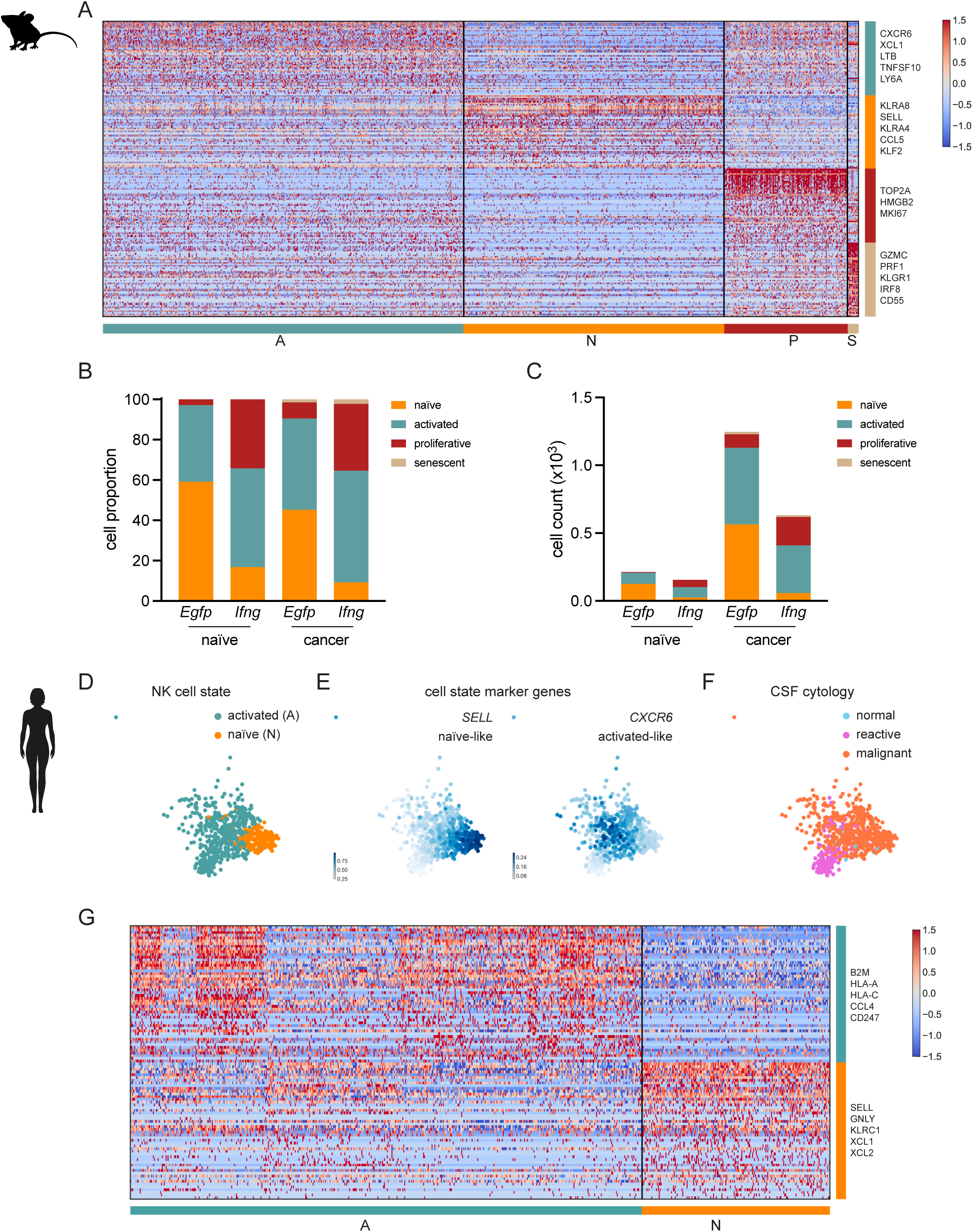
Characterization of leptomeningeal NK cells. (**A**) Heatmap showing scaled, zero-centered expression of top 50 genes *per* mouse NK cell state (one state vs. the rest; n = 2,247 cells total). Mouse NK cells were subsetted from ‘NK cell’ and ‘Proliferative T/NK cell’ clusters (Fig. 2B) based on the expression of Nk1.1 (cell surface) and *NKG7* (gene), and the lack of CD3 and TCRβ (cell surface). Naïve mouse NK cells are characterized based on single-cell RNA- and CITE-seq as CD62L^high^, activated NK cells are CD62L^low^, proliferative NK cells are CD62L^low^*MKI67*^+^, and senescent NK cells are CD55^+^ KLGR1^+^. See also Fig. 5 and Methods. (**B**) Proportion of NK cell states in naïve and metastasis-bearing, *Egfp*- or *Ifng*-overexpressing mice. (**C**) Cell counts of NK cell states in naïve and metastasis-bearing, *Egfp*- or *Ifng*-overexpressing mice. (**D**) UMAP showing NK cell states in human CSF (n = 763 cells); *NKG7*+ NK cell cluster from Fig. 1B was subsetted. (**E**) Projection of mouse naïve-like NK marker *SELL* (CD62L) and activated-like marker *CXCR6* onto human NK cells (MAGIC-imputed counts are plotted). (**F**) CSF cytology classification of human NK cells. (**G**) Heatmap showing scaled, zero-centered expression of top 50 genes *per* human NK cell state (one state vs. the rest).

**Fig. S20.**
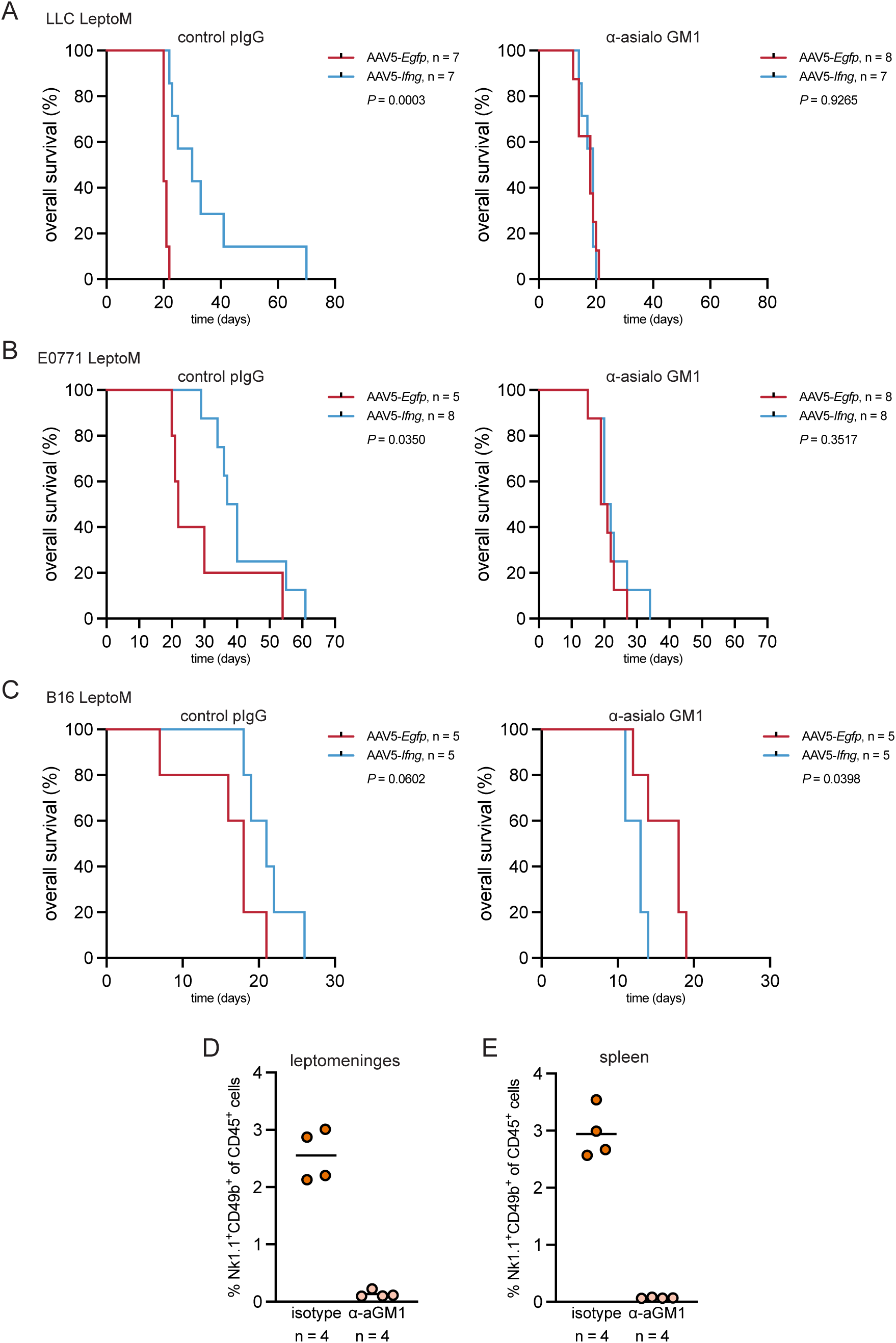
NK cells are the downstream cytotoxic effectors of leptomeningeal IFN-γ. (**A**) Kaplan-Meier plot showing survival of LLC LeptoM-bearing C57Bl/6-*Tyr*^c-2^ animals overexpressing *Egfp* or *Ifng* in the leptomeninges, depleted with control polyclonal antibody (left graph) or antibody targeting asialo-GM1 (logrank test). (**B**) Kaplan-Meier plot showing survival of E0771 LeptoM-bearing C57Bl/6-*Tyr*^c-2^ animals overexpressing *Egfp* or *Ifng* in the leptomeninges, depleted with control polyclonal antibody (left graph) or antibody targeting asialo-GM1 (logrank test). (**C**) Kaplan-Meier plot showing survival of B16 LeptoM-bearing C57Bl/6 animals overexpressing *Egfp* or *Ifng* in the leptomeninges, depleted with control polyclonal antibody (left graph) or antibody targeting asialo-GM1 in one experiment (logrank test). (**D**) Efficiency of systemic asialo-GM1-targeting depletion of NK cells in naïve C57Bl/6 animals, quantified in leptomeninges with flow cytometry. (**E**) Efficiency of systemic asialo-GM1-targeting depletion of NK cells in naïve C57Bl/6 animals, quantified in spleen with flow cytometry.

